# Why do eukaryotic proteins contain more intrinsically disordered regions?

**DOI:** 10.1101/270694

**Authors:** Walter Basile, Marco Salvatore, Claudio Bassot, Arne Elofsson

## Abstract

Intrinsic disorder is much more abundant in eukaryotic than in prokaryotic proteins. However, the reason behind this is unclear. It has been proposed that the disordered regions are functionally important for regulation in eukaryotes, but it has also been proposed that the difference is a result of lower selective pressure in eukaryotes. Almost all studies intrinsic disorder is predicted from the amino acid sequence of a protein. Therefore, there should exist an underlying difference in the amino acid distributions between eukaryotic and prokaryotic proteins causing the predicted difference in intrinsic disorder. To obtain a better understanding of why eukaryotic proteins contain more intrinsically disordered regions we compare proteins from complete eukaryotic and prokaryotic proteomes.

Here, we show that the difference in intrinsic disorder origin from differences in the linker regions. Eukaryotic proteins have more extended linker regions and, in particular, the eukaryotic linker regions are more disordered. The average eukaryotic protein is about 500 residues long; it contains 250 residues in linker regions, of which 80 are disordered. In comparison, prokaryotic proteins are about 350 residues long and only have 100-110 residues in linker regions, and less than 10 of these are intrinsically disordered.

Further, we show that there is no systematic increase in the frequency of disorder-promoting residues in eukaryotic linker regions. Instead, the difference in frequency of only three amino acids seems to lie behind the difference. The most significant difference is that eukaryotic linkers contain about 9% serine, while prokaryotic linkers have roughly 6.5%. Eukaryotic linkers also contain about 2% more proline and 2-3% fewer isoleucine residues. The reason why primarily these amino acids vary in frequency is not apparent, but it cannot be excluded that the difference is serine is related to the increased need for regulation through phosphorylation and that the proline difference is related to increase of eukaryotic specific repeats.

## Introduction

Eukaryotic proteomes have evolved to differ significantly from prokaryotic ones to deal with the increased complexity in a eukaryotic cell. The most notable differences are that (i) eukaryotic proteins are on average longer [1–4] (ii) multi-domain proteins are more abundant in eukaryotes [5–7], (iii) multicellular eukaryotic proteomes contain more domain repeats [8], and (iv) eukaryotic proteins have more intrinsically disordered regions [9]. Possible reasons for these differences include that eukaryotic cells have an increased need for regulation, that they contain organelles, the increased complexity of genes and genomes and a chaperone system enabling the folding of larger proteins [10].

The increased length of eukaryotic proteins is at least partly a consequence of them containing more domains that are arranged in a modular fashion. These have evolved mainly by fusion and terminal losses or gains of domains [11]. With a larger fraction of multi-domain proteins, it follows that eukaryotic proteins should have more linker regions - connecting the domains. Linker regions vary substantially in lengths and are often intrinsically disordered [12].

The increased number of repeats appears to be a feature of multicellular organisms [8]. Repeat proteins are common in signaling and are associated with some cancers. These repeats have been proposed to provide the eukaryotes with an extra source of variability to compensate for low generation rates [13].

The origin of the increase in intrinsically disordered regions in eukaryotic proteins is less well understood. Intrinsic disorder is frequent in all eukaryotic phyla, and even among viral proteins [14]. There is a spectrum of different types of disorder, spanning from increased flexibility to completely disordered proteins. Intrinsically disordered proteins exist among many classes of proteins with different functions, but the difference between eukaryotic and prokaryotic proteins appears to be maintained. On average fewer than 10% of the residues in prokaryotes are disordered regions compared with more than 20% in eukaryotes [15]. Disordered regions are overrepresented in regulatory proteins [16], providing a possible explanation for why the need for such regions is increased in eukaryotes.

It should, however, be remembered that the vast majority of studies of intrinsic disorder are based on predictions [17]. Intrinsic disorder predictions use amino acid frequencies and patterns found in the sequences. Although the best predictors use factors such as conservation and correlation between amino acid positions [18], even simple predictors that only use the amino acid frequency detects the difference between eukaryotes and prokaryotes [19]. Polar and charged amino acids, together with proline, are the most disorder-promoting residues. Thus, proteins with a higher fraction of these types of residues are (predicted to be) more disordered. Also, the average “disorder propensity”, as measured by the TOP-IDP scale [20], is significantly higher for eukaryotic proteins than for prokaryotic proteins.

For a protein family and over evolutionary time there are many possibilities for amino acid frequencies to change without the loss of function [21]. Most protein domain families contain members that have fewer than 20% sequence identities [22], i.e., at least 80% of the amino acids can be replaced. Protein design experiments have also shown that it is possible to design functional proteins with a limited, or biased, set of amino acids, which can be exemplified by some extreme cases such as the design of a protein without any charged residues [23] or SH3 domains with only a subset of amino acids [24]. Therefore, if it would exist a pressure to change the amino acid frequencies in the entire proteome, it should be possible for an organism to mostly adapt.

The general trend of amino acid gains and losses have been studied before, and it was proposed that the amino acids that appeared to increase in frequency (except serine) were not incorporated in the first genetic code [25]. However, the statistical methodology used in that study has been questioned [26]. The frequency of tyrosine has decreased in Metazoans compared to yeast [27], while histidine and serine frequencies increase from high-temperature thermophiles to prokaryotic mesophiles and further to eukaryotes while valine shows the opposite trend [28]. The trend of increasing polar amino acids in eukaryotes can be explained if the universal last common ancestor (LUCA) was oily, i.e., it was enriched in hydrophobic amino acids [29]. In this scenario, a selective pressure would be present to increase the number of polar amino acids, and thereby the predicted disorder. However, this model does not explain why this increase primarily occurred in eukaryotes and not in prokaryotes.

The GC content of a genome also influences the frequency of amino acids as the codons coding for different amino acids have a different fraction of GCs. It has been shown that genes encoded by high GC are more disordered [30] this trend is particularly strong among recently created genes but exist even among ancient genes [31]. Amino acids that have a high amount of GC among their codons are more frequent in genomes with high GC content, and amino acids whose codons have high GC content are more disorder-promoting than amino acid with low GC content. This difference explains why *de novo* created in yeast (low GC) are ordered while in Drosophila (high GC) these proteins are disordered. Peng et al. [30] proposed that the age of the amino acids could explain this trend. Old amino acids should then be disorder-promoting and use GC-rich codons while newer amino acids are mostly order-promoting and use GC-poor codons [31].

It has been proposed that intrinsic disorder, as well as the other features separating eukaryotic and prokaryotic proteins, is a result of low selective pressure and small effective population size [15]. The authors argue that low selective pressure in eukaryotes causes the expansion of non-coding regions in eukaryotic genomes, as there is no strong purifying selection to keep it compact. Supporting this argument is that young eukaryotic genes are on average more disordered than ancient ones [31]. However, there exist a large number of functionally important intrinsically disordered regions, see recent reviews [32, 33]. One of the functions for disordered regions in proteins is to present short linear motifs that are important for binding [16], further post-translational modification, in particular, phosphorylation, is a fundamental mechanism in the regulation of eukaryotic cell differentiation, as well as many other processes [34] and occurs preferentially in intrinsically disordered regions [35]. The functional importance indicates that disorder is not always a consequence of lower selection pressure, but also provides essential functions.

In this study, we ask what the molecular properties that underlie the difference in intrinsic disorder between eukaryotes and prokaryotes are. First, we show that the difference in disorder largely can be contributed to that linker regions in eukaryotes are not only more abundant but also more disordered. After that, we show that the difference in disorder can largely be attributed to a difference in serine, proline, and isoleucine frequencies.

## Material and Methods

### Datasets

The dataset used in this study originated from the complete bacterial, archaeal and eukaryotic proteomes from UniProt [36] as of December 2017. However, the frequency of some amino acids are strongly dependent on the GC content of the genome, see Figure S1 and in both prokaryotic kingdoms there exist a significant fraction of genomes with high GC content, see Figure S2. Differences in GC composition complicates the comparison of amino acid distributions. We examined several alternative methods to take the effect of the difference of amino acid frequencies caused by the difference in GC into account, including ANOVA test, see Table S1, and projections into an average GC content (data not shown), with similar results. However, we do believe that the easiest way to compensate for GC is to ignore all genomes with high GC content. This removal makes it possible to compare trends both between proteomes and within protein families without compensating for a difference in GC. Therefore we excluded all genomes with a GC content of more than 60% or less than 20% from this study. The resulting set of genomes have a similar GC content in all three kingdoms (43-44% GC with a standard deviation of 8%), see Figure S2b.

This final dataset contains 26,274,724 protein sequences from 6,373 genomes. The proteomes are divided into 4,905 bacterial, 308 archaeal, and 975 eukaryotic. Here, all species from the following taxa, Mycoplasma, Spiroplasma, Ureaplasma, and Mesoplasma were ignored as they have another codon usage - which influences the expected amino acid frequencies.

As differences at a proteome level can be due both to a different number of proteins of a particular type and differences within these type of proteins, we divided the complete proteomes into subgroups. We first identified the set of protein domain families from Pfam [37, 38] that are present in both bacterial and eukaryotic genomes. We retained all Pfam domains present in at least ten species from two distinct kingdoms among the annotated “full” alignments in Pfam, and where none of the kingdoms make up of more than 99.9% of all the proteins, resultingin a set of 4,165 “shared” Pfam domain families, out of these 1,764 are common to all three kingdoms.

We define “Shared proteins” as all proteins that contain at least one of these domains. Proteins that only contain the Pfam domains that are specific to a kingdom are referred to as (kingdom) specific proteins, and proteins without any Pfam domains are referred to as No Domain proteins, see Figure 1.

**Figure 1.**
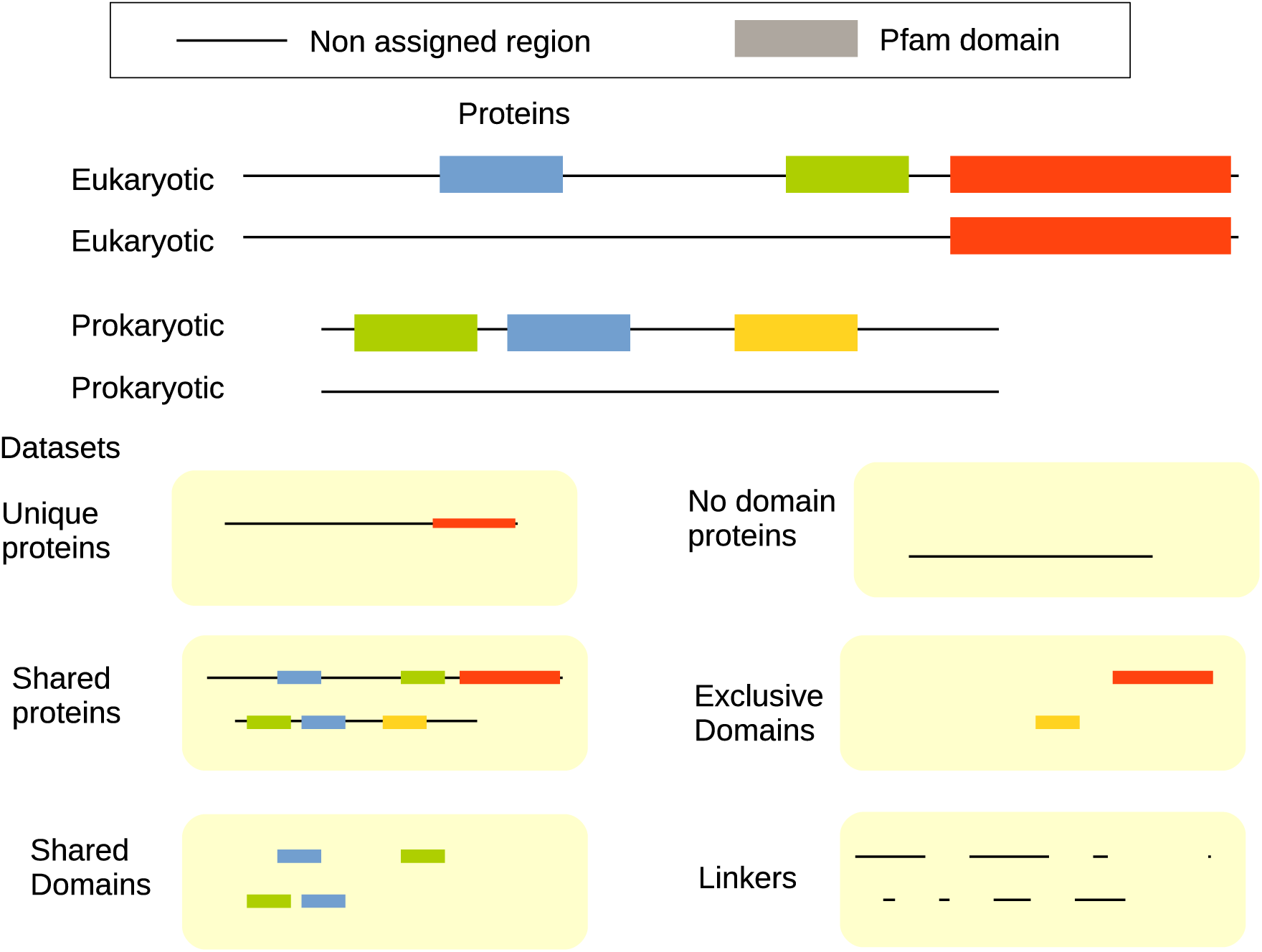
Representation of how the proteins are divided into six subsets in this study. All the proteins from all the proteomes in the two kingdoms are first divided into three groups. The ones that only contain unique domains to one of the kingdoms, the ones without any domains and the set that contains at least one shared domain. The last set is then divided into three regions: Shared Domains, Exclusive Domains, and Linkers. Note that a protein can contain several domains of each type, e.g., it can contain more than one shared domain.

Even within one group of proteins differences might be due to an abundance of different regions or differences within regions. Therefore, we divided the proteins the proteins which shared domains into regions, see Figure 1. The regions of each protein corresponding to any of the 4,165 Pfam domains that exist both in prokaryotes and eukaryotes are referred to as “Shared domains”. All regions assigned to any other Pfam domain are “Specific domains”; and all residues that are not assigned to a Pfam domain are classified to be within a “Linker region”.

The entire proteomes are thereby divided into six sets. First, the proteins are subdivided into three subgroups: kingdom specific proteins, no domain proteins and shared proteins. Next, the shared proteins are divided into three regions, shared domains, kingdom specific domains, and linker regions.

For each of these six groups, we analyzed length, disorder, amino acid frequencies, and other properties independently. Here, it should be noted that we report the average number of amino acids in a set and not the average length of each region, i.e., if a protein contains two shared domains both domains are used to calculate the length. As shown in Figure 1 a protein can contain zero, one or multiple regions of a particular type.

The processed datasets, as well as all scripts, are available from a figshare repository [39].

### Disorder prediction

For each protein, we estimated the intrinsic disorder by using two tools: IUPred [19] and the TOP-IDP scale [20]. IUPred exploits the idea that in disordered regions, amino acid residues form less energetically favorable contacts than the ones in ordered regions. IUPred does not rely on any external information besides the amino acid sequence, and for this reason is extremely fast and suitable to predict disorder for a large dataset. We used the IUPred to predict long disordered regions. For each protein, we used the default cut-off and assigned a residue to be disordered if its IUPred value is greater than 0.5. Using the short version of IUPred gives almost identical results (data not shown), i.e., the results are robust when it comes to the choice of disorder predictor.

The TOP-IDP scale [20], assigns a disorder propensity score to each amino acid, and it is based on statistics on previously published scales. For each protein region, a TOP-IDP value is calculated as the average of the TOP-IDP values of all its residues.

## Results

First, we compare the average length and disorder content for proteins in the different domains of life. The proteins are divided into three groups and three regions using Pfam, see Figure 1. We choose to use Pfam even if sometimes the exact definitions of individual Pfam domains can be discussed [40].

In total, the proteomes contain 26M proteins. About half (14M) of the protein belong to the group *shared proteins*, i.e., they contain at least one Pfam domain that exists in at least two kingdoms. Thiese proteins can, therefore, be considered to be ancient. The 4M *kingdom specific proteins* only contain Pfam domains that are unique to one of the three kingdoms. These proteins are most likely more recent inventions and could perform functions specific to properties unique to one of the kingdoms. Finally, we have in total 8.3M proteins without any annotated Pfam domain, most likely these are the youngest proteins, but could also incorporate fast evolving proteins. Next, we also studied the 14M proteins with a shared domain in more details by dividing these into three regions: regions with a shared Pfam domain, regions with a kingdom specific Pfam domain and linker regions.

### Longer linkers make eukaryotic proteins longer

As has been documented many times before, eukaryotic proteins are longer and more disordered than prokaryotic proteins, see Figure 2 and Tables S2-S4. Protein with shared domains are on average longer than proteins without such domains, and proteins without domains are even shorter, see Figure 2. However, in all three groups of proteins, the Eukaryotic proteins are longer. It has been proposed that multi-domain proteins are almost twice as frequent in eukaryotes [4]. However, this cannot be the only explanation of the difference in length as even proteins without any assigned domains are longer in eukaryotes.

**Figure 2.**
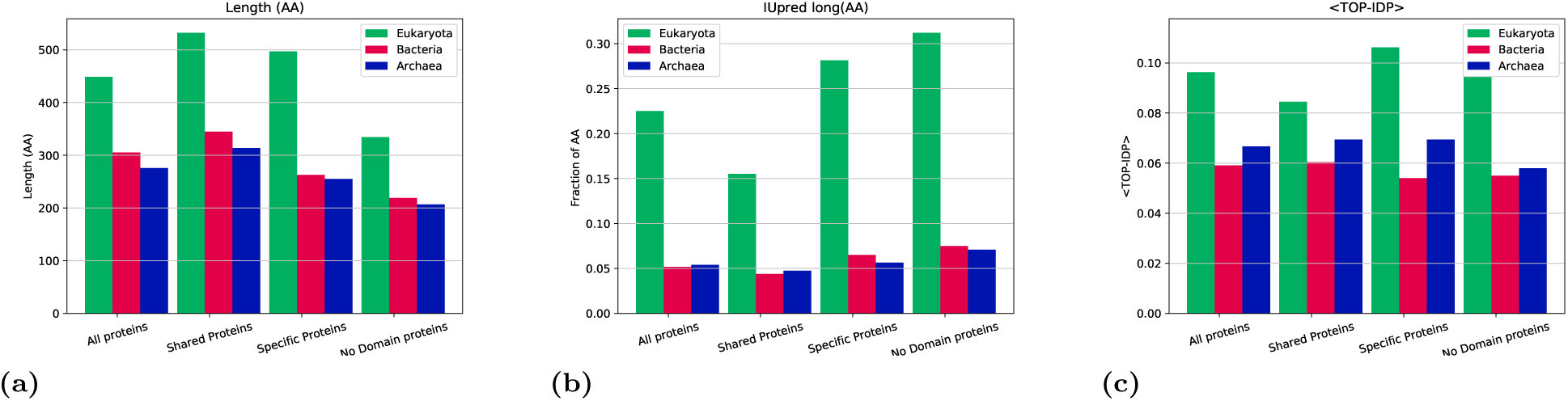
Average properties of proteins from di_erent kingdoms; (a) average length, (b) fraction of residues predicted to be disordered by IUPred (c) average TOP-IDP scores.

It would be possible that the linker regions contain conserved domains not present in Pfam. Although it is certain that some domains are not yet present in Pfam and that some distant members of a domain family can be missed we do not believe this to be a significant difference between eukaryotes and prokaryotes when Pfam is based on thousands of complete genomes from all kingdoms of life. Therefore, our earlier explanation (more multi-domain proteins in eukaryotes) to the length difference is quite likely to be incorrect [4]. This proposal can be contributed to the fact that we in that earlier study assumed that long linker regions contained unassigned domains, which is likely not always the case.

To understand the origin of the difference in length we choose to study the proteins that contain a Pfam domain that exist in two kingdoms in more detail. Among the 14 million proteins with shared domains the average length of eukaryotic proteins is 536 residues vs. 361 for bacterial protein and 337 for archaeal proteins, see Figure 2 and Tables S2-S4. The number of residues in the shared domains is roughly equal in the three kingdoms, 218 to 233 residues. The average number of residues assigned to kingdom-specific domains is 27 in bacteria, 19 in archaea, and 49 in eukaryotes, i.e., the absolute number of residues assigned to these unique domains do only marginally contribute to the difference in average lengths, see Figure 3. The number of residues in linker regions, 258 in eukaryotes vs. 113 residues in bacteria and 106 in archaea, thus cause *>* 80% of the length difference. In eukaryotes, 48% of all residues are assigned to linkers, while in prokaryotes only 31%.

**Figure 3.**
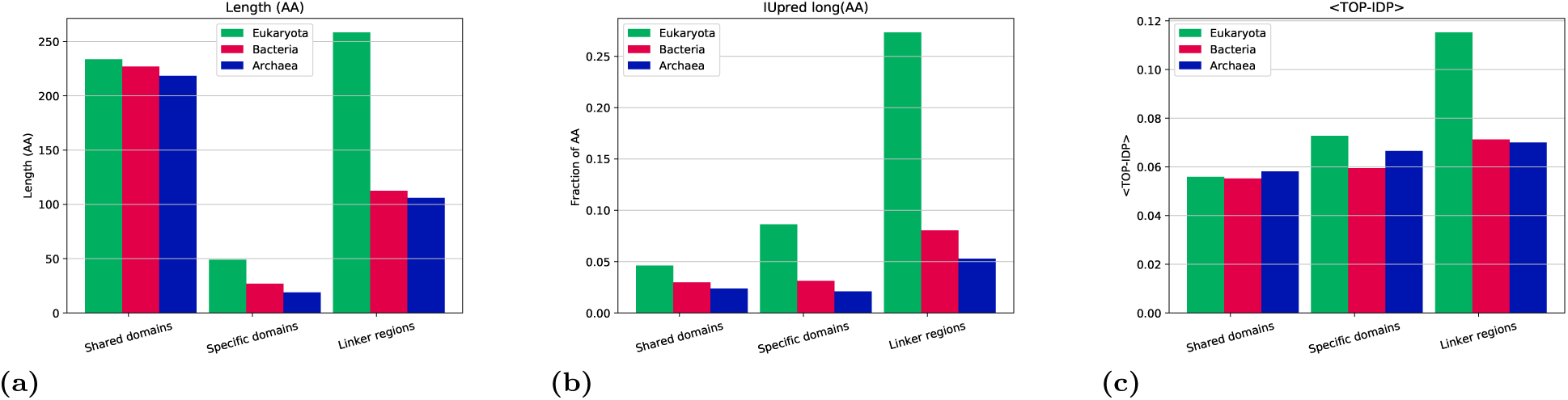
Average properties of regions from different kingdoms; (a) average length, (b) average TOP-IDP; scores and fraction of residues predicted to be disordered by (c) IUPred-short and (d) IUPred-long.

Linkers can be located at one of the termini or between two domains. In all three kingdoms, roughly 40% (35-43%) of the linker residues are located at each of the termini, and fewer residues (from 18% in archaea to 27% in eukaryotes) are located between domains (central). Linkers at all locations are on average more than twice as long in eukaryotes than in prokaryotes.

### Eukaryotic linkers are more disordered

Eukaryotic proteins are more disordered than prokaryotic ones, see Figure 2b and c. In agreement with earlier studies [15, 41–43] 5% of the residues in prokaryotes are predicted to be disordered compared with 23% in eukaryotes. The increase in disorder is independent of the type of protein, but proteins that are specific to eukaryotes are more disordered than the ones that contain a shared domain. Eukaryotic proteins without any Pfam domains are most disordered with 31% disordered residues, supporting earlier observations that young proteins are more disordered than old ones, at least in some eukaryotes [31].

The shared domains contain least disordered residues in all kingdoms (fewer than 5%), see Figure 3. The kingdom specific domains are more disordered than shared domains in eukaryotes (9%) but not among the prokaryotes. In all three kingdoms, the linker regions are the most disordered regions, as reported before [17]. However, the major difference is that eukaryotic linker regions are more disordered than prokaryotic linkers, 27% vs. 5-8%. Independent on location the eukaryotic linkers are more disordered, see Table S2-S4. The difference in disorder can therefore not only be contributed to that linker regions are more abundant in eukaryotic proteins, but these regions are also more disordered in eukaryotes.

### Eukaryotic proteins have 100 disordered residues and prokaryotic 15

On average eukaryotic proteins are roughly 450 residues long and contain about 23% disordered residues, while prokaryotic proteins are 300 residues long and contain 5% disorder residues. This results in that on average a eukaryotic protein contains roughly 100 disordered residues, while in average prokaryotic protein only have 15 disordered residues, see Figure 4. Proteins with eukaryotic specific domains contain more disordered residues (139) than the ones with shared domains (83) although they are shorter. The group of proteins without domains have the highest fraction of intrinsically disordered residues but is the shortest group; therefore the number of disordered residues is similar to the average (104 vs. 101). All types of prokaryotic proteins contain roughly the same number of disordered residues (13 to 15).

**Figure 4.**
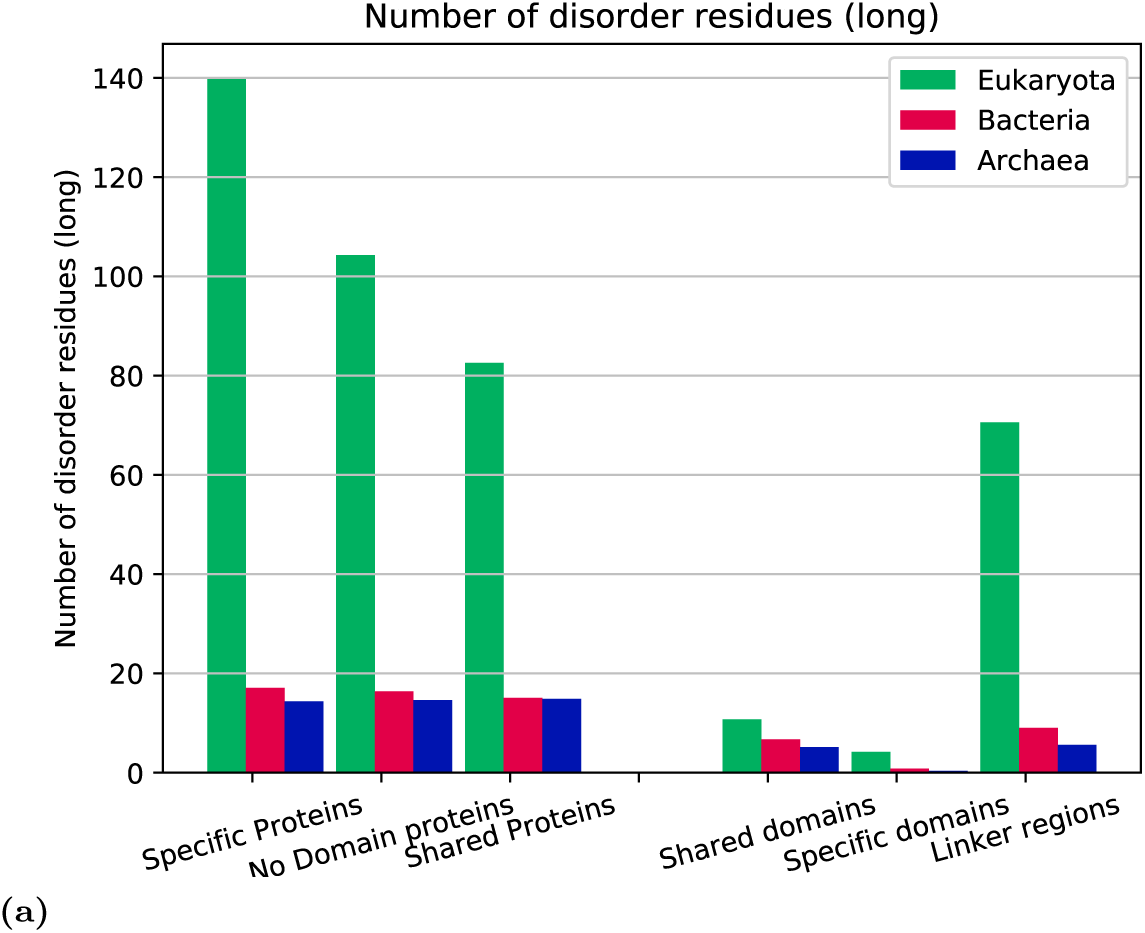
Average number of residues predicted to be disordered in different protein groups and regions.

In summary, eukaryotic proteins are 150 residues longer than prokaryotic proteins. This difference is mainly due to more abundant linker regions, both at the termini and in between domains. The linker regions in eukaryotes are more disordered than prokaryotic linker regions. A typical eukaryotic protein contains 100 disordered residues and 90 of these residues are located in linker regions, see Table S2. In contrast, a bacterial protein contains on average 15 disordered residues, and 9 of these are located in linker regions, see Table S3.

### AA frequencies in eukaryotic linkers are unique

Above we show that eukaryotic proteins are more disordered due to that their linker regions are both longer and more disordered. The difference in (predicted) intrinsic disorder is primarily due to differences in amino acid frequencies. Therefore next, we studied the difference in amino acid frequencies between linker regions in eukaryotes and prokaryotes. For comparison, we compared differences between the shared domains.

The different regions can be clustered using the similarity in amino acid frequency, see Figure 5. Here, it is clear that eukaryotic linker region is an outlier, clearly separated from all other regions. Next, it can also be observed that all regions in archaea cluster together, while the remaining eukaryotic domains and bacterial regions form a third cluster. Figure 5 highlights that eukaryotic linker regions has evolved to become quite distinct from all other regions and might indicate that there exist specific selective preferences acting on these linkers.

**Figure 5.**
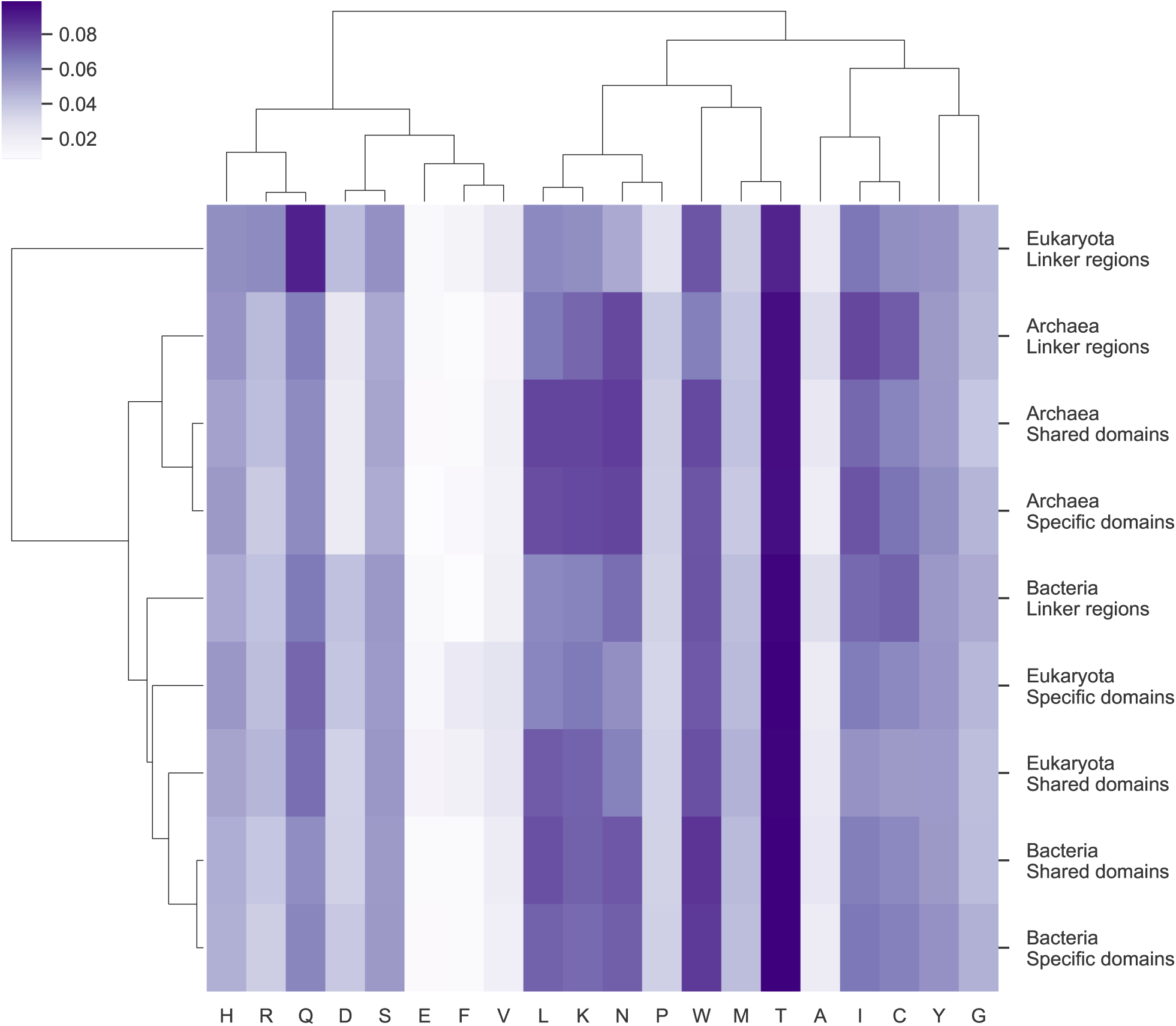
Heat map showing the similarity of amino acid frequency profiles in different regions as measured by the Pearson correlation coefficient. The color of each cell represents the Pearson correlation, according to the reference color bar.

In Figure 6 and Table 1 it can be seen that: (a) the difference between linker regions is higher than between shared domains (0.9% vs. 0.5% on average), confirming the clustering in Figure 5 and (b) that there exist three amino acids (isoleucine, serine and proline) whose frequencies differ by more than 1.5% between eukaryotic linkers and linkers in both prokaryotes. Also, there is one amino acid (glutamine) where the frequency in archaeal linkers is more than 1.5% lower in archaeal than in eukaryotic linkers. However, the frequency of glutamine in bacterial linkers is similar to the frequency in eukaryotic linkers. Therefore, it seems as if there is some specific reason for the low glutamine frequency in archaea. As confirmed by the ANOVA test these three amino acids are the most significant differences between the eukaryotic and bacterial linkers, see Table S1.

**Table 1.** Amino acid frequencies for each protein region and kingdom.

|  | Eukaryota |  |  | Bacteria |  |  | Archaea |  |  |
| --- | --- | --- | --- | --- | --- | --- | --- | --- | --- |
|  | Shared domains | Specific domains | Linker regions | Shared domains | Specific domains | Linker regions | Shared domains | Specific domains | Linker regions |
| TRP | 1.5% | 1.3% | 1.1% | 1.0% | 1.0% | 1.1% | 1.0% | 0.8% | 1.1% |
| PHE | 4.6% | 4.3% | 3.6% | 4.3% | 4.2% | 4.2% | 4.0% | 3.8% | 3.9% |
| TYR | 3.4% | 3.3% | 2.7% | 3.4% | 3.5% | 3.4% | 3.6% | 3.6% | 3.8% |
| ILE | 6.3% | 5.8% | 4.9% | 7.5% | 7.2% | 6.9% | 8.1% | 7.9% | 7.8% |
| MET | 2.3% | 2.1% | 2.2% | 2.4% | 2.1% | 2.9% | 2.3% | 2.0% | 3.0% |
| LEU | 9.8% | 9.8% | 8.8% | 9.8% | 9.9% | 9.7% | 9.5% | 9.5% | 9.5% |
| VAL | 7.1% | 6.5% | 5.8% | 7.2% | 7.0% | 6.2% | 7.9% | 7.8% | 7.1% |
| ASN | 4.2% | 4.4% | 4.5% | 4.2% | 4.6% | 4.8% | 3.9% | 4.5% | 4.4% |
| CYS | 1.8% | 2.2% | 1.5% | 1.0% | 1.0% | 0.8% | 1.1% | 1.2% | 0.9% |
| THR | 5.5% | 5.4% | 5.7% | 5.3% | 5.4% | 5.5% | 5.0% | 4.8% | 4.9% |
| ALA | 7.6% | 7.4% | 7.5% | 8.4% | 8.2% | 7.5% | 7.8% | 7.5% | 6.4% |
| GLY | 7.3% | 6.1% | 6.0% | 7.6% | 7.2% | 6.0% | 7.9% | 7.7% | 6.5% |
| ARG | 5.0% | 5.5% | 5.8% | 4.7% | 4.6% | 4.9% | 5.1% | 5.4% | 5.6% |
| ASP | 5.4% | 5.5% | 5.6% | 5.4% | 5.7% | 5.4% | 5.4% | 5.8% | 5.4% |
| HIS | 2.5% | 2.6% | 2.4% | 2.1% | 1.9% | 1.8% | 1.8% | 1.7% | 1.6% |
| GLN | 3.4% | 3.9% | 4.2% | 3.5% | 3.8% | 4.0% | 2.2% | 2.2% | 2.4% |
| SER | 6.8% | 7.1% | 8.9% | 5.9% | 6.1% | 6.5% | 5.9% | 5.9% | 6.3% |
| LYS | 5.3% | 6.0% | 5.8% | 6.0% | 6.3% | 7.2% | 6.2% | 6.7% | 7.3% |
| GLU | 5.7% | 6.5% | 6.7% | 6.4% | 6.6% | 7.0% | 7.0% | 7.5% | 7.9% |
| PRO | 4.4% | 4.2% | 5.9% | 3.9% | 3.6% | 4.0% | 4.2% | 3.7% | 4.3% |

**Figure 6.**
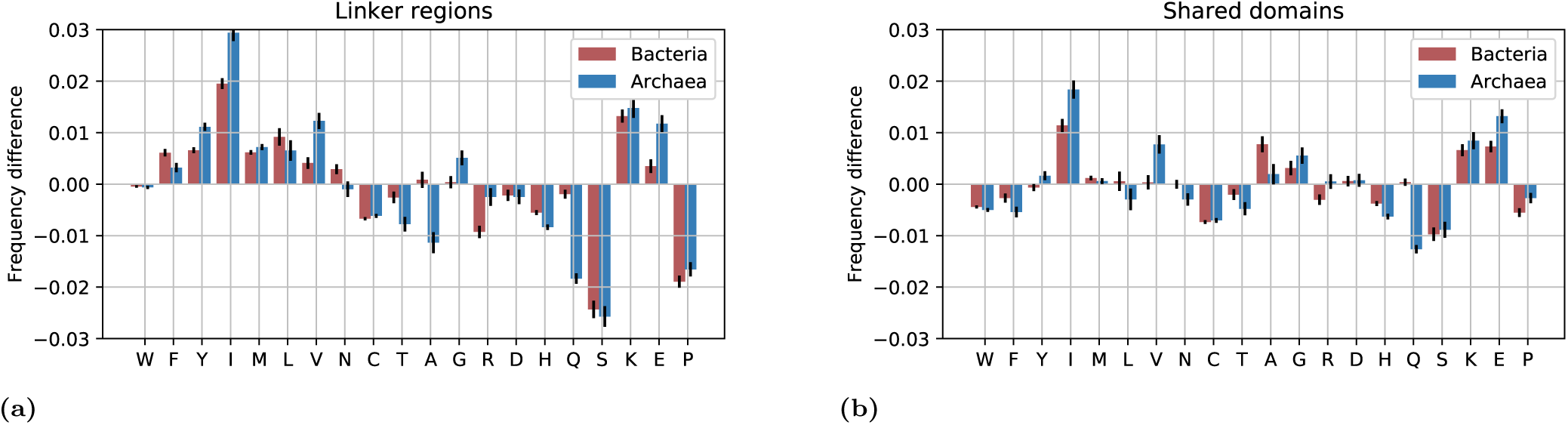
Amino acid frequency between eukaryotes and prokaryotes for linker regions.

From Figure 6 it is clear that in hydrophobic residues are less frequent in eukaryotic linkers, while some disorder-promoting (polar) residues are increased and some decreased. The two disorder-promoting residues, lysine, and glutamic acid, are less frequent in eukaryotic linkers and glutamine is equally frequent in eukaryotic an bacterial linkers. In contrast, as highlighted above, proline and serine are more frequent in the eukaryotic linkers.

Within the shared domains the differences in amino acid frequencies are in general smaller, and there is no systematic decrease of hydrophobic amino acids in eukaryotic proteins. It can be noted that the three amino acids with most substantial differences in frequency (isoleucine, serine, and proline) are shifted in the same direction both in domains and in linkers but that the differences are smaller within the domains, possibly indicating the existing of to some system-wide preferential difference of these amino acids in the different kingdoms.

Further, from the clustering of amino acids in Figure 5 it can be seen that that amino acids can be divided into three groups. The first group (RSPQTWCH) is enriched in eukaryotic proteins. These amino acids are mainly polar. one subgroup (RSPQT) is most abundant in eukaryotic linkers, see Table 1, while the rest (WCH) are more abundant in eukaryotic domains.

The second cluster (VGYIAFL) consist of mainly hydrophobic amino acids. These are all rare in all linkers, but in particular in eukaryotic linkers, and one subcluster (VGYI) is enriched in archaea and the other (AFL) in bacteria.

The third cluster contains the remaining polar amino acids (KEND). Lysine and glutamine are enhanced primarily in the prokaryotic linker regions, but also in the archaeal domains, while asparagine and aspartic acid are evenly distributed in all regions.

In summary, it does not appear to be as simple as that disorder is increased in eukaryotic linkers. Instead, there are mainly three amino acids whose frequencies differ between eukaryotic and prokaryotic linkers. These amino acids (isoleucine, serine, and proline) contribute to an increased disorder in eukaryotic linkers. If these three amino acids are ignored there is no difference in disorder propensity, see Figure S3 Taken all this into account it is not clear if the increased disorder is primarily a consequence of changes in amino acids frequencies, or if the increased disorder drives the difference in amino acid frequencies, a classic chicken and egg problem.

### The difference in amino acid frequencies is widespread

Next, we studied the frequency of individual amino acids in different phyla, see Figure 7 and S4. Serine is increased in the linkers of all eukaryotic phyla. For proline and isoleucine, the trend seems to be consistent, but the frequency differences are not observed in alveolata, and there exist a few bacterial phyla (Chloroflexi, Planctomycete, Actinobacteria, and Acidobacteria) that have frequencies similar to the what is observed in eukaryotes. These are all phyla with high GC content, see Figure S5. The codons for proline are enriched in GC while the isoleucine codons are GC poor. Therefore, the genomic GC levels could explain the frequencies in these phyla. It could be considered that the average GC content also could explain the observations in Alveolata, which has a low GC content, but the GC content in Metazoa and Viridiaeplanta are similar to Alveolate, see Figure S5, and they have typical eukaryotic amino acid frequencies. Therefore, we do believe that there is some other unknown explanation why proline frequencies are low, and isoleucine frequencies are high in Alveolata.

**Figure 7.**
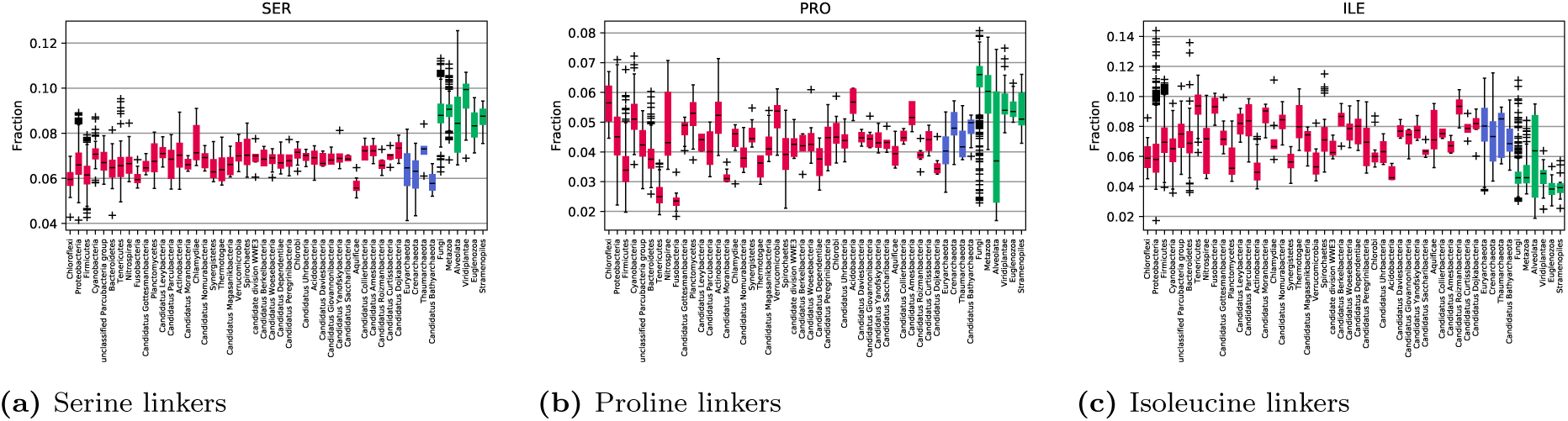
Frequency shifts of serine, proline, and isoleucine in proteomes grouped by phylum. Bacterial groups are red, eukaryotic green, and archaeal blue.

That fact that the serine frequency is increased in all eukaryotic phyla indicates that the differences are most likely an acient feature. Further supporting this, we examined the amount of serine in the complete proteome, i.e., not only linkers, in the prokaryotic phylum that has been proposed to bridge the gap between prokaryotes and eukaryotes [44]: the archaea Lokiarchaeota contain 6.4% serine, while one of the most primitive eukaryotes, *Giardia lamblia* contains 9.2% serine, indicating that the increase of serine is something that occurred early after LUCA [45] and has been maintained in all eukaryotic lineages ever since. Proline and isoleucine frequencies also support the old ancestry as the eukaryotes *Giardia lamblia* contain typical eukaryotic levels of proline (4.7%) and isoleucine (5.7%), while the archaea Lokiarchaeota contains less proline (3.5%) and more isoleucine (10.2%).

### Frequency differs in most protein families

Eukaryotic proteomes are in general larger than prokaryotic proteomes; this is partly due to an expansion of protein families. For functional reasons, different protein families also have different amino acid distributions. Therefore, it is possible that the differences in the amino acid frequency that we observe when studying an entire proteome are due to the different frequencies of families with different amino acid distributions. Therefore, we examined the amino acid frequency of all shared Pfam domains. The reason to study domains and not the linkers is that the linkers are difficult to align and differs significantly in length, while the domains are of similar length and already aligned in Pfam.

In Figure 8 the differences between the amino acid frequencies in the prokaryotic domains are compared with the amino acid frequencies in the corresponding eukaryotic domains. Only Pfam families with at least 100 members in a bacteria and eukaryotes are included to avoid statistical outliers (archaea was ignored in this filtering). In most families the trends from the linkers are conserved; In 84% of the families, the eukaryotic members have more serine, in 80% fewer isoleucine and in 70% more proline. We tried to identify any trends among the families with high or low frequencies, both by examining individual families and by mapping onto

**Figure 8.**
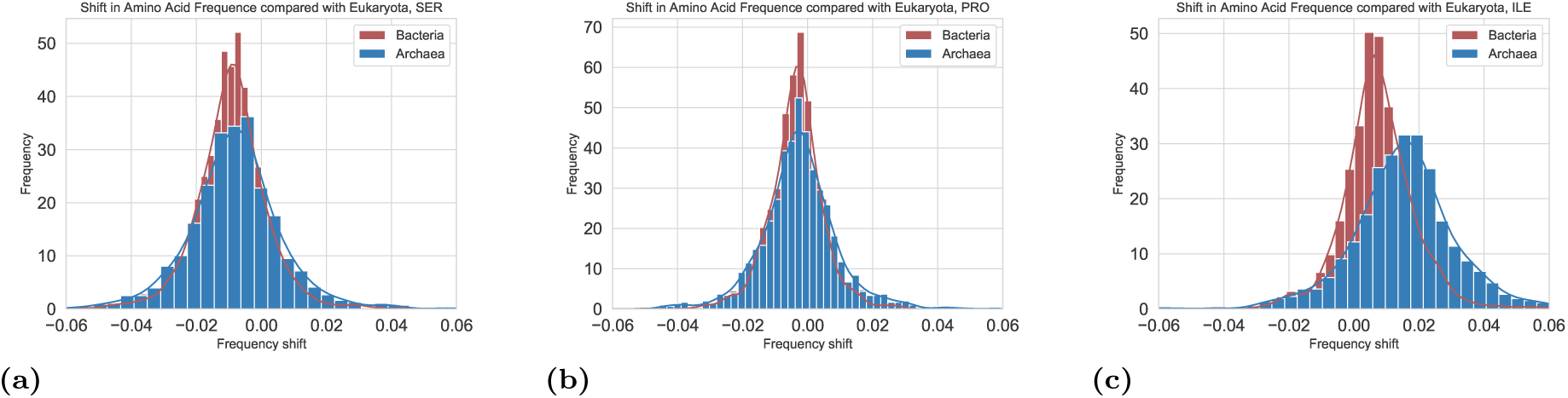
Distribution of differences of amino acids frequencies in protein with the same Pfam family. Only Pfam families that contain at least 100 members in two kingdoms are included in the comparison.

GO-terms using GO-slim. The tool pfam2go [46] was used to obtain GO terms [47, 48] for each family. The GO terms were then mapped to the GOslim generic category. The GO terms with largest differences in amino acid frequency are listed in Tables S7-**??**. To the best of our ability, we cannot identify any particular subset of proteins where the difference in frequency significantly differs from the general picture.

Another difference between eukaryotes and prokaryotes is that eukaryotic cells have different organelles. The amino acid content of proteins in different organelles differs; therefore it would not be implausible that the different amino acid frequencies could be affected by the compartmentalization of the eukaryotic cell. However, in all membrane and non-membrane parts of all organelles the frequencies of serine and proline are higher in eukaryotes than in prokaryotes, see Table S5, and in all, except three (vacuole plus mitochondrial and plastid membrane) organelles the isoleucine frequency is lower in eukaryotes.

Some bacteria in the Planctomycetes, Verrucomicrobiae, and Chlamydiae bacterial superphylum have quite complex membranes possibly indicating primitive organelles [49]. However, all these phyla have levels of these amino acids for bacteria, see Figure 7, i.e., the compartmentalization of the eukaryotic cell does not seem to explain the differences in frequencies.

### Serine is enriched in disorder regions

Next, we studied if the difference in amino frequency in different structural regions of a protein. For this, we used the shared domains again, as the structural information of the linkers is very limited. We only used the Pfam families where there was at least one structure available in PDB and assumed that the secondary structure was conserved within the Pfam family.

Using the secondary structure annotation, available from Pfam, we then assigned each protein into four categories, Helix, Sheet or Coil given the most frequent annotation among all structures in that column in the Pfam multiple sequence alignment. Positions that were unassigned, i.e., corresponding to the parts of the Pfam domains that are not present in any PDB structure, we do refer to as disordered. There might be many other reasons why the region is not present in the PDB, but this definition is related to the missing residues, which has been used for training many disorder predictors. As expected the serine and proline frequency is highest in loops and disordered regions, while isoleucine frequency is lowest in these regions, see Figure 9.

**Figure 9.**
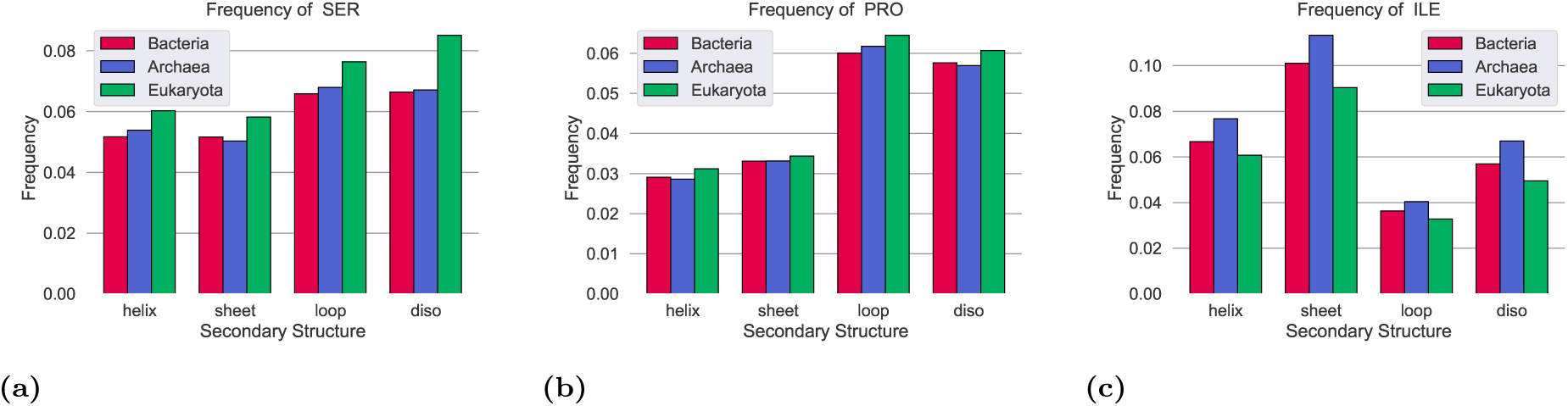
Frequency of serine in different secondary structures in bacterial and eukaryotic proteins. Here only proteins from genomes with 40-50% GC are used.

The difference in amino acid frequency between the kingdoms exist in all secondary structures, but the increase of serine is largest in disordered regions where it is close to 2% higher in eukaryotes than in prokaryotes.

## Discussion

Above we note that eukaryotic linkers are longer and more disordered than prokaryotic linkers regions. We further showed that this is primarily due to a shift in the frequency of three amino acids, serine and proline are more frequent in eukaryotic linkers while isoleucine is less frequent. Here, we will speculate about possible explanations for these differences.

### Lower selective pressure in eukaryotic linkers

Functional selection is typically considered to be the dominant force shaping proteome evolution, but it cannot be excluded that secondary effects such as the cost of producing an amino acid or codon preferences also affect the general trend of amino acid frequencies [50].

The population size of eukaryotes is in general smaller than for prokaryotes causing a lower selective pressure. Further, the amount of intrinsic disorder is lower than expected by chance in both eukaryotes and prokaryotes [31]. Therefore it is possible that the lower selective pressure could explain why eukaryotes contain more disordered residues [15]. The basis of this argument is that the majority of the disordered residues are unfavorable, at least in prokaryotes, and therefore a selective pressure act against them. However, this is not the case, such as arginine, some disorder-promoting residues are less frequent than expected by chance, while others, including lysine, are more frequent, see figure S6.

However, we show that the difference in disorder is mainly due to the differences of frequencies of only three amino acids and is unique to the linker regions, indicating that the simple explanation of a lower selective pressure cannot be the full story. Further, in at least in some eukaryotic proteins it has been shown that disorder has a clear functional role [32, 33], leading to the question: what phenotypic aspects of eukaryotic linker regions explain the enrichment of serine and proline and the decrease of isoleucine.

In bacteria, it has been proposed that one reason to reduce the frequency of an amino acid is the energetic cost to produce it [51]. Serine is among the least costly amino acids to make both aerobically and anaerobically [51, 52], see Table S6. Proline is also cheaper than most amino acids to make, while isoleucine is among the most expensive ones. Therefore, the cost of producing the amino acids could not explain the differences in frequency.

### Serine is a target for kinases

One possible reason for the higher fraction of serine in eukaryotic organisms is that serine, together with threonine, are targets for ser/thr kinases [53]. Phosphorylation of serine and threonine is one the critical regulatory pathways in eukaryotes, but also present in archaea [54]. It is also established that phosphorylation frequently occurs in intrinsically disordered sites [55].

Phosphorylation can occur at three amino acids, in addition to serine, threonine, and tyrosine. Threonine and tyrosine frequencies show no increase in eukaryotic linker regions, see Figure S4. If phosphorylation by kinases is the primary reason for the serine frequency difference between eukaryotes and prokaryotes one can wonder why only serine frequency is increased. However, it might be due to that about 75% of the known targets for kinases are serine [56]. However, it might also be related to the fact that serine is disorder-promoting while threonine and tyrosine are not.

Ser/thr kinases are prevalent in eukaryotes, but also exist in bacteria such as planctomycetes [57]. In the only fully sequenced genome of this phylum (*Planctomycetes bacterium GWA2 40 7*) there are only 6.1% serine. Further, the major family of ser/thr kinases, Pfam family Stk19 (PF10494), only exists in eukaryotes and in Halanaerobiales. In the 2783 Halanaerobiales sequences in UniProt [36] there is 5.8% serine, typical of a prokaryote.

It can be seen that serine and proline are less frequent than expected in prokaryotic linkers, see figure 10, indicating a selective pressure to reduce the amount of serine. The exact reason for this is unknown, but it has been reported that high serine levels are toxic [58].

**Figure 10.**
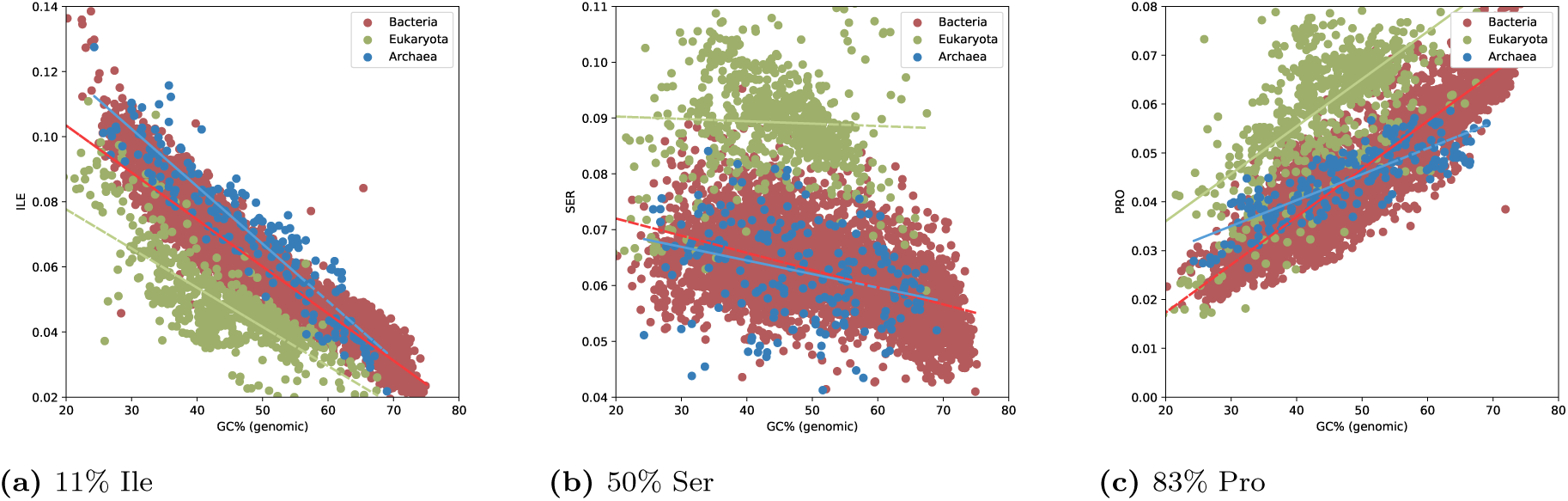
Frequency of isoleucine, serine, and proline vs. GC of the linker regions in the genomes. The amino acids are sorted after the GC content of their codons. The number represents the fraction of GC among the codons. The black line represents the expected frequency from codon usage only.

### Proline

In addition to being enriched in loops, proline is often unique to repeat proteins [59], and in particular, PPP and PPG repeats are more common in multicellular organisms [60]. Proline repeats are also often found in disordered regions that are important for binding in eukaryotic specific proteins such as P53 [61]. Proline is also a common amino acid in linker regions connecting domains [62]. As both repeats and multi-domain proteins are more frequent in eukaryotes, all these factors might contribute to the increase of proline in eukaryotic linkers.

### Isoleucine

Prokaryotes (but not eukaryotes), use a specific purine-rich sequence on the 5’ side to distinguish initiator AUGs from internal ones. The codons for isoleucine contains four adenosine (out of nine nucleotides), this could potentially contribute to the higher fraction of isoleucine in prokaryotes. However, as the differences between eukaryotes and prokaryotes also exist in C-terminal regions this can not be the only explanation for the difference, i.e., it is not apparent why isoleucine and not other hydrophobic residues are underrepresented in eukaryotic linkers.

### Conclusion

Here, we show that eukaryotic proteins are more disordered than prokaryotic proteins because they have more extended linker regions and that these linker regions are more disordered. Further, the difference in disorder content of these regions does not origin from a general increase of disorder-promoting residues; instead, it is a consequence of systematic shifts in the frequency of only three amino acids, serine, proline, and isoleucine. Serine and proline are more frequent in eukaryotic linkers than in prokaryotic once, while isoleucine is less frequent. These differences hold over all phyla and the clear majority of protein families. For serine, in particular, the difference can even be seen in all regions of proteins but is most pronounced in disordered regions. Are these shifts in amino acids frequencies primarily a consequence of increased intrinsic disorder in eukaryotic linkers or the reason behind this increase ? However, the observation that not all disorder-promoting amino acids are increased in eukaryotic proteins makes it clear that earlier explanations of the increased disorder in eukaryotic proteins is simplified.

## Acknowledgements

This work was supported by grants from the Swedish Research Council (VR-NT 2016-03798 to AE) and Swedish e-Science Research Center. The Swedish National Infrastructure provided computational resources for Computing (SNIC)

We do thank Nir Ben-Tal, Erich Bornberg-Bauer, David Liberles, Claudio Bassot and Marco Salvatore for valuable inputs and discussions.

## Supporting Information

**Table S1.** ANOVA F-test for contribution of different amino acids to the difference between eukaryotic and bacterial proteomes when including the GC genomic content.

| AA | F |
| --- | --- |
| Ser | 3058.361533 |
| Prp | 2487.525547 |
| Ile | 2173.393746 |
| Cys | 1169.086554 |
| His | 749.223044 |
| Arg | 510.144299 |
| Tyr | 489.430463 |
| Met | 450.230289 |
| Lys | 410.439484 |
| Phe | 364.499096 |
| Leu | 293.245268 |
| Val | 238.185813 |
| Gln | 180.921957 |
| Gly | 156.656403 |
| Ala | 142.546529 |
| Asn | 81.479070 |
| Thr | 60.226780 |
| Gly | 42.153414 |
| Asp | 37.907222 |
| Trp | 22.751912 |

**Table S2.** Summary of features for the different sets of proteins and protein regions in Eukaryota

| Eukaryota | Shared domains | Specific domains | Linker regions | N-terminal Linker regions | Central Linker regions | C-terminal Linker regions | All proteins | No Do-main proteins | Specific Proteins | Shared Proteins |
| --- | --- | --- | --- | --- | --- | --- | --- | --- | --- | --- |
| IUpred long(AA) | 0.0460 | 0.0862 | 0.2732 | 0.2896 | 0.2336 | 0.2798 | 0.2251 | 0.312 | 0.2814 | 0.1552 |
| IUpred short(AA) | 0.0385 | 0.0634 | 0.2589 | 0.2867 | 0.1996 | 0.2668 | 0.1984 | 0.2687 | 0.2398 | 0.1437 |
| <SEG> | 0.0346 | 0.0404 | 0.0993 | 0.1056 | 0.0859 | 0.1009 | 0.0869 | 0.1118 | 0.1006 | 0.0669 |
| <TOP-IDP> | 0.0559 | 0.0728 | 0.1152 | 0.1139 | 0.1176 | 0.1149 | 0.0963 | 0.112 | 0.1063 | 0.0846 |
| <Hydrophobicity> (Hessa) | 0.9741 | 1.0405 | 1.1249 | 1.1112 | 1.1327 | 1.1342 | 1.0806 | 1.1143 | 1.1134 | 1.0498 |
| Length (AA) | 233.6505 | 49.0138 | 258.3515 | 98.3297 | 68.5544 | 91.4674 | 448.7465 | 334.2222 | 496.8121 | 532.2085 |
| Number of disorder residues (long) | 10.7536 | 4.2259 | 70.5803 | 28.4728 | 16.0142 | 25.5971 | 101.0064 | 104.2736 | 139.8248 | 82.5723 |
| Number of disorder residue (short) | 8.9977 | 3.1088 | 66.8898 | 28.192 | 13.6812 | 24.401 | 89.0183 | 89.8144 | 119.157 | 76.4827 |
| Low complexity residues | 8.0786 | 1.9791 | 25.6432 | 10.3822 | 5.8906 | 9.2285 | 39.0099 | 37.3726 | 49.9907 | 35.5892 |
| TRP | 0.0148 | 0.0131 | 0.0114 | 0.0107 | 0.0106 | 0.0127 | 0.0123 | 0.0115 | 0.0114 | 0.013 |
| PHE | 0.0457 | 0.043 | 0.0359 | 0.0364 | 0.0352 | 0.0357 | 0.0391 | 0.0361 | 0.038 | 0.0409 |
| TYR | 0.0343 | 0.0328 | 0.0272 | 0.0267 | 0.0272 | 0.0278 | 0.0297 | 0.0274 | 0.0286 | 0.0311 |
| ILE | 0.0629 | 0.0576 | 0.0489 | 0.0482 | 0.0505 | 0.0485 | 0.0535 | 0.0496 | 0.0507 | 0.0565 |
| MET | 0.0226 | 0.0215 | 0.0224 | 0.0258 | 0.0198 | 0.0206 | 0.0223 | 0.0219 | 0.0223 | 0.0225 |
| LEU | 0.0978 | 0.0984 | 0.0883 | 0.0898 | 0.088 | 0.0871 | 0.0923 | 0.0893 | 0.0934 | 0.0932 |
| VAL | 0.0714 | 0.0655 | 0.0583 | 0.0566 | 0.0596 | 0.0593 | 0.0611 | 0.0573 | 0.0574 | 0.0649 |
| ASN | 0.042 | 0.044 | 0.0448 | 0.0445 | 0.0456 | 0.0444 | 0.0448 | 0.0454 | 0.0448 | 0.044 |
| CYS | 0.0177 | 0.022 | 0.0152 | 0.0149 | 0.0152 | 0.0154 | 0.0166 | 0.0174 | 0.0161 | 0.0165 |
| THR | 0.0552 | 0.0537 | 0.0572 | 0.0584 | 0.0579 | 0.0554 | 0.0569 | 0.0589 | 0.0561 | 0.0562 |
| ALA | 0.0762 | 0.0741 | 0.0748 | 0.0759 | 0.0741 | 0.0742 | 0.0744 | 0.0735 | 0.074 | 0.0751 |
| GLY | 0.0734 | 0.0613 | 0.0602 | 0.0564 | 0.0622 | 0.0632 | 0.0615 | 0.0576 | 0.0557 | 0.0666 |
| ARG | 0.0505 | 0.0552 | 0.0583 | 0.0587 | 0.0558 | 0.0594 | 0.0563 | 0.0596 | 0.058 | 0.0542 |
| ASP | 0.0535 | 0.0555 | 0.0564 | 0.0549 | 0.0587 | 0.0566 | 0.0548 | 0.0536 | 0.0557 | 0.055 |
| HIS | 0.0247 | 0.0259 | 0.0239 | 0.0246 | 0.0233 | 0.0236 | 0.0244 | 0.0259 | 0.0237 | 0.024 |
| GLN | 0.0343 | 0.0392 | 0.0423 | 0.0421 | 0.0425 | 0.0427 | 0.0413 | 0.0436 | 0.0449 | 0.0382 |
| SER | 0.0684 | 0.0705 | 0.0895 | 0.0943 | 0.0862 | 0.0863 | 0.0837 | 0.0915 | 0.0869 | 0.0781 |
| LYS | 0.0535 | 0.0599 | 0.0582 | 0.0563 | 0.059 | 0.06 | 0.0572 | 0.0566 | 0.0591 | 0.0569 |
| GLU | 0.0566 | 0.0646 | 0.0665 | 0.0623 | 0.0689 | 0.0696 | 0.0639 | 0.0648 | 0.0681 | 0.0619 |
| PRO | 0.0443 | 0.0420 | 0.0594 | 0.0621 | 0.0591 | 0.0564 | 0.0536 | 0.0581 | 0.0544 | 0.0510 |
| <Alpha propensity> | -0.0102 | -0.0043 | -0.0123 | -0.0127 | -0.0129 | -0.0112 | -0.0105 | -0.012 | -0.0081 | -0.0109 |
| <Beta propensity> | -0.0316 | -0.0363 | -0.0528 | -0.0521 | -0.0537 | -0.0528 | -0.0455 | -0.0502 | -0.0489 | -0.0418 |
| <Coil propensity> | -0.0141 | -0.0158 | -0.009 | -0.0085 | -0.0088 | -0.0099 | -0.0111 | -0.0092 | -0.0115 | -0.0119 |
| <Turn propensity> | -0.0702 | -0.0646 | -0.0482 | -0.0494 | -0.0476 | -0.0473 | -0.0554 | -0.0495 | -0.0523 | -0.0596 |

**Table S3.** Summary of features for the different sets of proteins and protein regions in Bacteria

| Bacteria | Shared domains | Specific domains | Linker regions | N-terminal Linker regions | Central Linker regions | C-terminal Linker regions | All proteins | No Do-main proteins | Specific Proteins | Shared Proteins |
| --- | --- | --- | --- | --- | --- | --- | --- | --- | --- | --- |
| IUpred long(AA) | 0.0297 | 0.0312 | 0.0804 | 0.0729 | 0.0800 | 0.0873 | 0.0517 | 0.0748 | 0.065 | 0.0438 |
| IUpred short(AA) | 0.0264 | 0.0261 | 0.1290 | 0.1498 | 0.0616 | 0.1492 | 0.0656 | 0.0911 | 0.0767 | 0.0578 |
| <SEG> | 0.0363 | 0.0363 | 0.0574 | 0.0676 | 0.0509 | 0.0506 | 0.0473 | 0.0606 | 0.0574 | 0.0429 |
| <TOP-IDP> | 0.0553 | 0.0595 | 0.0712 | 0.0514 | 0.0905 | 0.0798 | 0.0591 | 0.055 | 0.054 | 0.0604 |
| <Hydrophobicity> (Hessa) | 0.9738 | 1.0018 | 1.051 | 0.9865 | 1.0995 | 1.0876 | 0.9979 | 1.0045 | 0.9926 | 0.9967 |
| Length (AA) | 226.8661 | 26.8114 | 112.4711 | 43.8507 | 26.6907 | 41.9297 | 305.2008 | 218.9736 | 262.699 | 344.5751 |
| Number of disorder residues (long) | 6.7446 | 0.8353 | 9.0414 | 3.1963 | 2.1358 | 3.6588 | 15.7684 | 16.3861 | 17.0764 | 15.0894 |
| Number of disorder residue (short) | 5.9927 | 0.699 | 14.5046 | 6.5709 | 1.6434 | 6.2563 | 20.0298 | 19.9507 | 20.1384 | 19.9064 |
| Low complexity residues | 8.2408 | 0.9728 | 6.456 | 2.9635 | 1.3584 | 2.1201 | 14.4427 | 13.2703 | 15.0686 | 14.7837 |
| TRP | 0.0104 | 0.0099 | 0.011 | 0.0113 | 0.009 | 0.0118 | 0.0111 | 0.0128 | 0.0122 | 0.0105 |
| PHE | 0.0432 | 0.042 | 0.0421 | 0.0458 | 0.0382 | 0.0408 | 0.0445 | 0.0476 | 0.0465 | 0.0436 |
| TYR | 0.0338 | 0.0349 | 0.0339 | 0.0335 | 0.032 | 0.0355 | 0.0343 | 0.0375 | 0.0358 | 0.0334 |
| ILE | 0.0746 | 0.0725 | 0.0687 | 0.0713 | 0.0666 | 0.0673 | 0.073 | 0.0711 | 0.0724 | 0.0737 |
| MET | 0.0237 | 0.0212 | 0.0285 | 0.0417 | 0.0198 | 0.0205 | 0.0244 | 0.0236 | 0.0236 | 0.0247 |
| LEU | 0.0985 | 0.0987 | 0.0975 | 0.1016 | 0.0931 | 0.096 | 0.0987 | 0.0985 | 0.1005 | 0.0987 |
| VAL | 0.0716 | 0.0696 | 0.0622 | 0.061 | 0.0646 | 0.0619 | 0.0676 | 0.0634 | 0.0666 | 0.0687 |
| ASN | 0.0424 | 0.0463 | 0.0483 | 0.048 | 0.0481 | 0.0486 | 0.0449 | 0.0492 | 0.048 | 0.0434 |
| CYS | 0.0103 | 0.0104 | 0.0085 | 0.0087 | 0.0072 | 0.009 | 0.0097 | 0.0107 | 0.0082 | 0.0096 |
| THR | 0.0532 | 0.0538 | 0.0546 | 0.0563 | 0.0556 | 0.0519 | 0.0538 | 0.0554 | 0.0555 | 0.0531 |
| ALA | 0.0836 | 0.0822 | 0.0752 | 0.0733 | 0.0785 | 0.0751 | 0.079 | 0.0747 | 0.0765 | 0.0802 |
| GLY | 0.0762 | 0.0721 | 0.0602 | 0.0562 | 0.0639 | 0.0619 | 0.0696 | 0.064 | 0.066 | 0.0712 |
| ARG | 0.0472 | 0.0465 | 0.0487 | 0.0457 | 0.0494 | 0.0515 | 0.0478 | 0.0457 | 0.0461 | 0.0485 |
| ASP | 0.0541 | 0.0569 | 0.0543 | 0.049 | 0.0606 | 0.0558 | 0.0534 | 0.0534 | 0.0525 | 0.0534 |
| HIS | 0.0209 | 0.0191 | 0.0184 | 0.0172 | 0.0187 | 0.0194 | 0.0192 | 0.0178 | 0.0168 | 0.0199 |
| GLN | 0.0347 | 0.0378 | 0.0405 | 0.0394 | 0.0418 | 0.0408 | 0.0362 | 0.0377 | 0.0388 | 0.0355 |
| SER | 0.0587 | 0.0611 | 0.0652 | 0.0677 | 0.0642 | 0.063 | 0.0623 | 0.0673 | 0.0664 | 0.0606 |
| LYS | 0.0605 | 0.0631 | 0.0721 | 0.0713 | 0.0681 | 0.0755 | 0.0656 | 0.0667 | 0.0653 | 0.0657 |
| GLU | 0.0639 | 0.0658 | 0.07 | 0.0618 | 0.0773 | 0.0742 | 0.0654 | 0.0640 | 0.0640 | 0.0660 |
| PRO | 0.0387 | 0.0361 | 0.0403 | 0.0394 | 0.0432 | 0.0394 | 0.0394 | 0.0389 | 0.0382 | 0.0395 |
| <Alpha propensity> | -0.005 | -0.004 | -0.001 | 0.0004 | -0.0031 | -0.0009 | -0.0039 | -0.0046 | -0.0041 | -0.0036 |
| <Beta propensity> | -0.0315 | -0.0334 | -0.0372 | -0.0295 | -0.0447 | -0.0406 | -0.0328 | -0.0323 | -0.0311 | -0.0329 |
| <Coil propensity> | -0.0175 | -0.0174 | -0.0178 | -0.0192 | -0.0159 | -0.0175 | -0.0175 | -0.0167 | -0.0174 | -0.0179 |
| <Turn propensity> | -0.0764 | -0.0729 | -0.0702 | -0.0795 | -0.0633 | -0.0651 | -0.0744 | -0.0723 | -0.0745 | -0.075 |

**Table S4.**
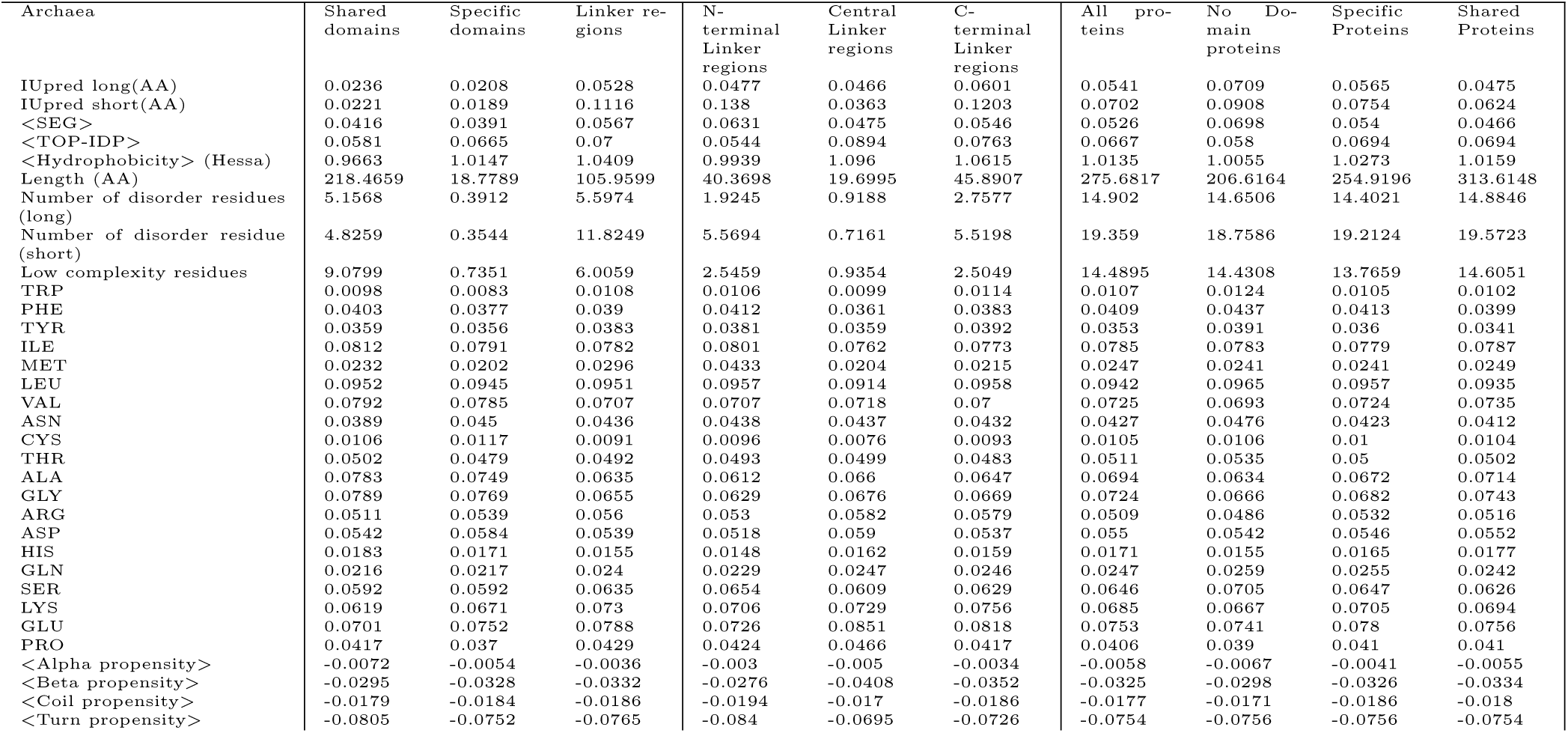
Summary of features for the different sets of proteins and protein regions in Archaea

**Table S5.**
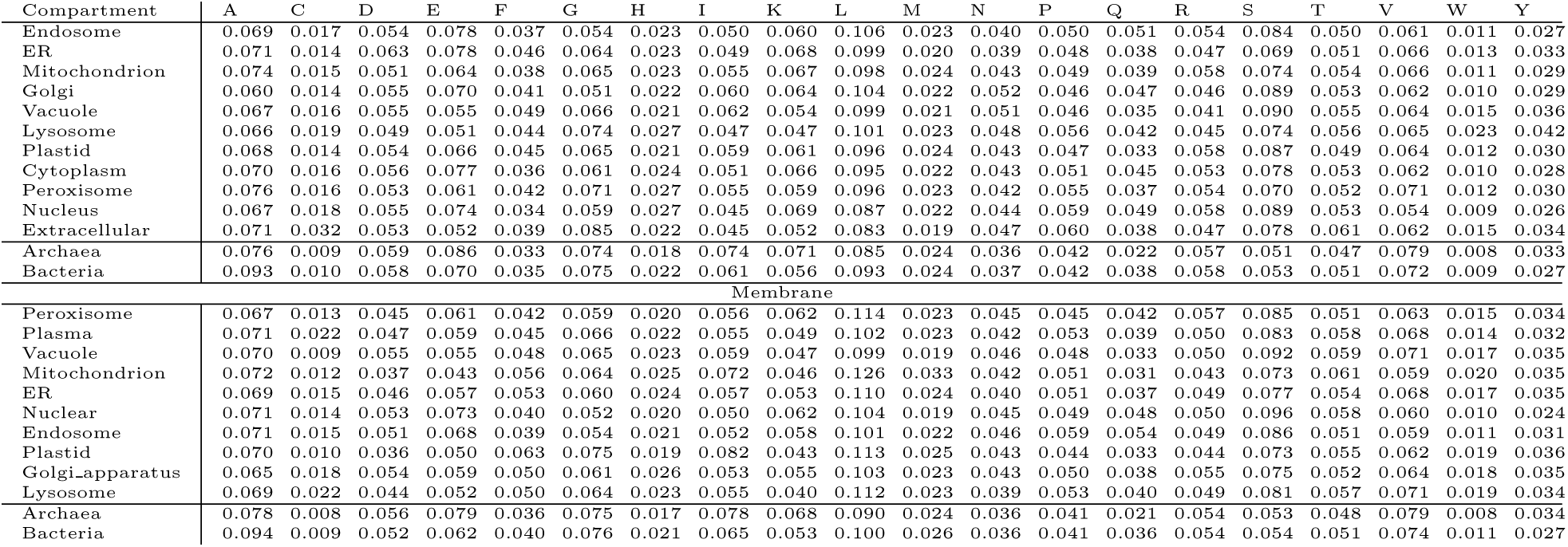
Frequency of amino acids in the different subcellular compartments. The sequences are taken from all Swissprot proteins with subcellular annotations.

**Table S6.** Anearobic cost and aerobic costs to make an amino acid. acids. Data in column one is from Akashi et al [51] and in column two and three from Raiford et al [52]. Units are the number of *PO*_4_ molecules. The table is sorted according to the bost in from the first study.

| AA | Aerobic [51] | Aerobic [52] | Anaerobic [52] |
| --- | --- | --- | --- |
| Gly | 11.7 | 14.5 | 1.0 |
| Ser | 11.7 | 14.5 | 1.0 |
| Ala | 11.7 | 14.5 | 2.0 |
| Asp | 12.7 | 15.5 | 3.0 |
| Asn | 14.7 | 18.5 | 6.0 |
| Glu | 15.3 | 9.5 | 2.0 |
| Gln | 16.3 | 10.5 | 3.0 |
| Thr | 18.7 | 21.5 | 9.0 |
| Pro | 20.3 | 14.5 | 7.0 |
| Val | 23.3 | 29.0 | 4.0 |
| Cys | 24.7 | 26.5 | 13.0 |
| Arg | 27.3 | 20.5 | 13.0 |
| Leu | 27.3 | 37.0 | 4.0 |
| Lys | 30.3 | 36.0 | 12.0 |
| Ile | 32.3 | 38.0 | 14.0 |
| Met | 34.3 | 36.5 | 24.0 |
| His | 38.3 | 29.0 | 5.0 |
| Tyr | 50.0 | 59.0 | 8.0 |
| Phe | 52.0 | 61.0 | 10.0 |
| Trp | 74.3 | 75.5 | 14.0 |

**Table S7.**
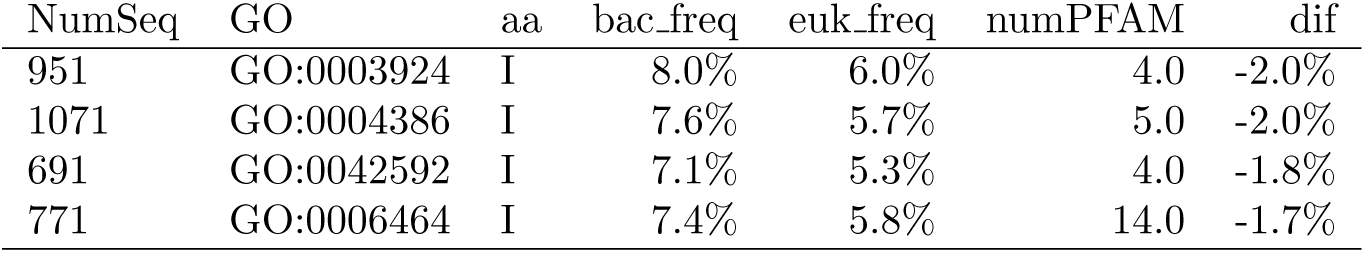
GO terms that differ with isoleoucine frequency differs with more than 1.5% betwen eukaryotes and bacteria. NumSeq is the minimum number of sequences in a Pfam family, numpFAM is that number of Pfam families with this GO term. The Fraction of families with decreased eukaryotic frequency: 77.9%

**Table S8.**
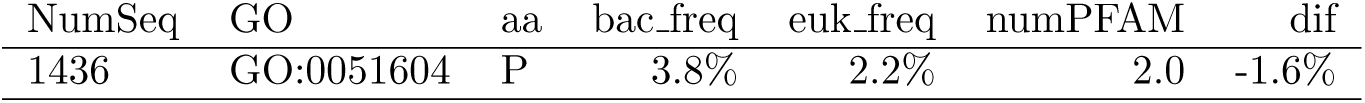
GO terms that differ with proline frequency differs with more than 1.5% betwen eukaryotes and bacteria. NumSeq is the minimum number of sequences in a Pfam family, numpFAM is that number of Pfam families with this GO term. The Fraction of families with increased eukaryotic frequency: 80.9 %

**Table S9.** GO terms that differ with serine frequency differs with more than 1.5% betwen eukaryotes and bacteria. NumSeq is the minimum number of sequences in a Pfam family, numpFAM is that number of Pfam families with this GO term. The Fraction of families with increased eukaryotic frequency: 86.6 %

| NumSeq | GO | aa | bac_freq | euk_freq | numPFAM | dif |
| --- | --- | --- | --- | --- | --- | --- |
| 1539 | GO:0005694 | S | 5.1% | 6.7% | 4.0 | 1.5% |
| 959 | GO:0003924 | S | 5.3% | 7.1% | 4.0 | 1.9% |
| 339 | GO:0008135 | S | 7.0% | 4.3% | 3.0 | -2.7% |
| 1439 | GO:0051604 | S | 7.8% | 6.3% | 2.0 | -1.5% |

**Figure S1.**
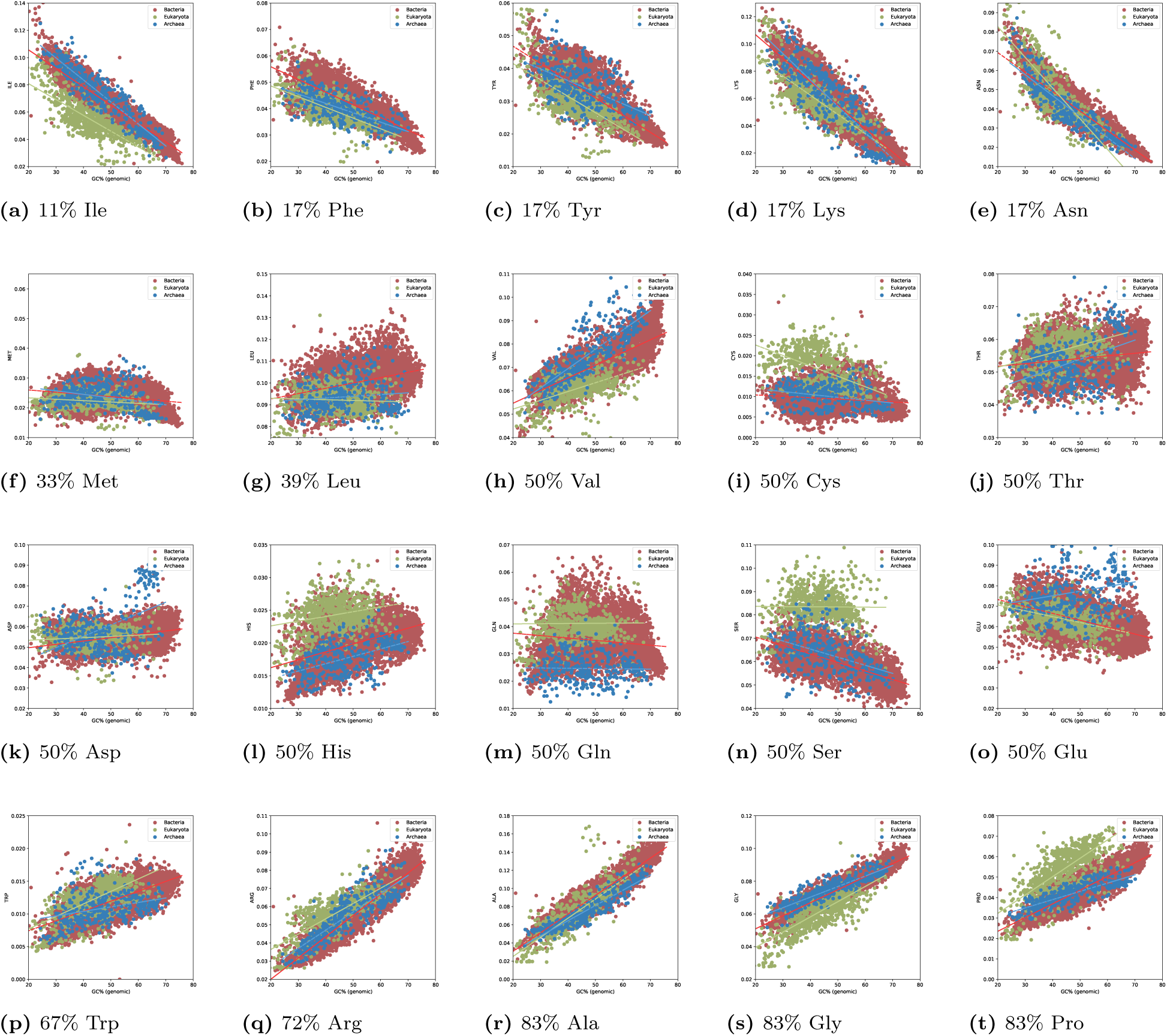
Frequency of all amino acids vs. GC of the genomes amino acids. The amino acids are sorted after the GC content of their codons. The number represents the fraction of GC among the codons. The straight lines represents linear fits for each kingdom independently.

**Figure S2.**
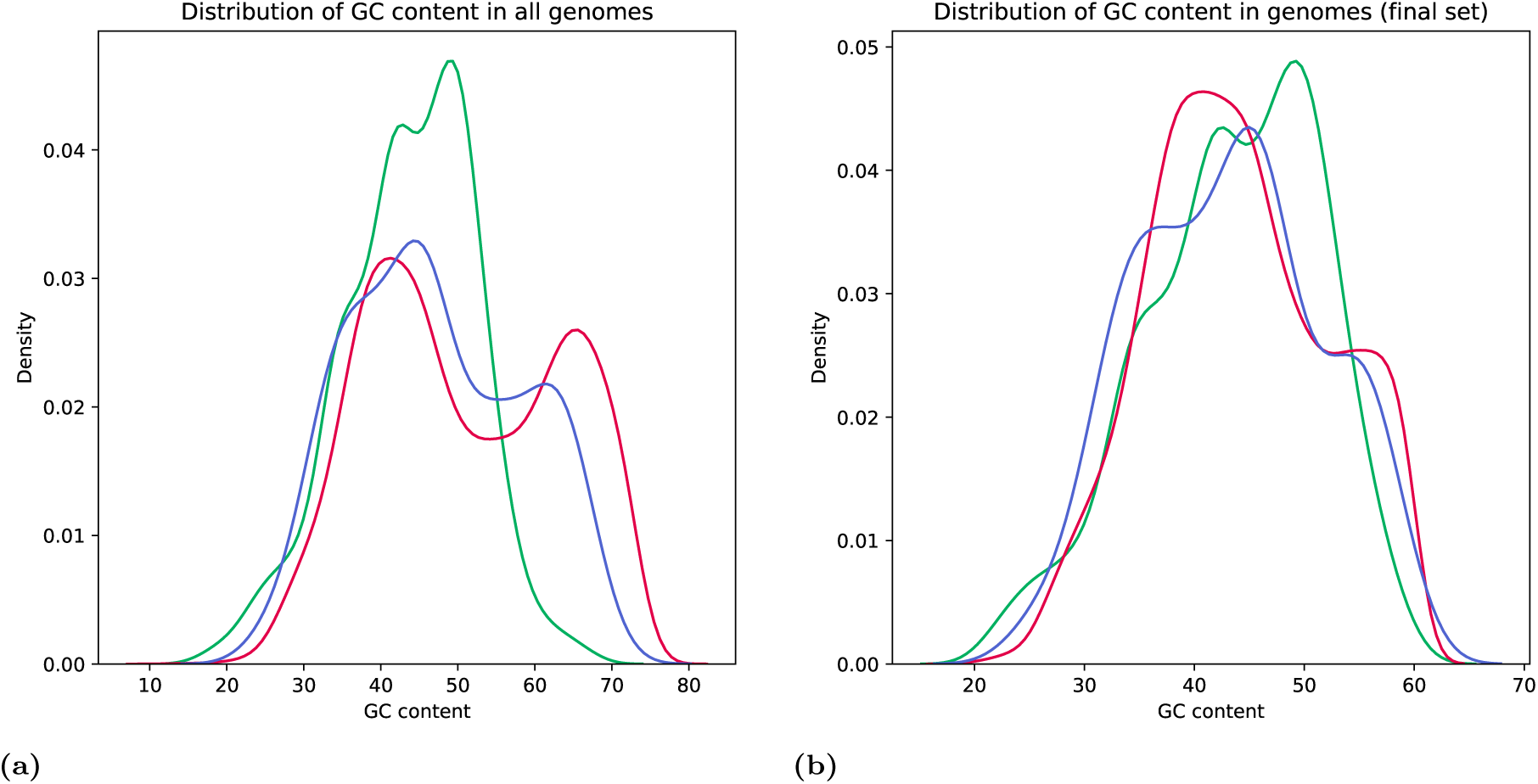
Density plot of the distribution of GC in genomes from different kingdoms. In a) all genome are shown and in b) only the genomes that remained after filtering for GC between 20 and 60%. When all genomes are present the average GC content of eukaryotes is 43.8%, and 51.0% for bacteria and 47.2% for archaea. After filtering the average GC is similar in all three kingdoms (43.2 to 44.0%) as are the standard deviations (8.0 to 8.4). Only 2.6% of the eukaryotic genomes are excluded (25 out of 1001), but 20% of the archaeal (75 out of 383) and 30% of the bacterial ones (2219 out of 7124).

**Figure S3.**
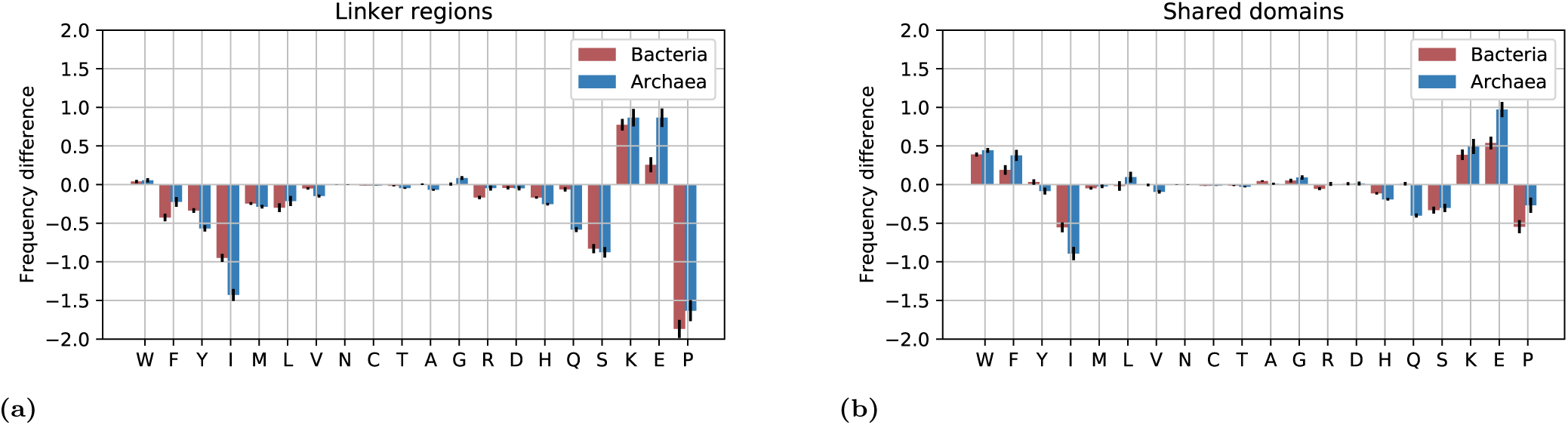
Difference in disorder propensity contributed by differences in amino acid frequency between linkers in the three kingdoms.

**Figure S4.**
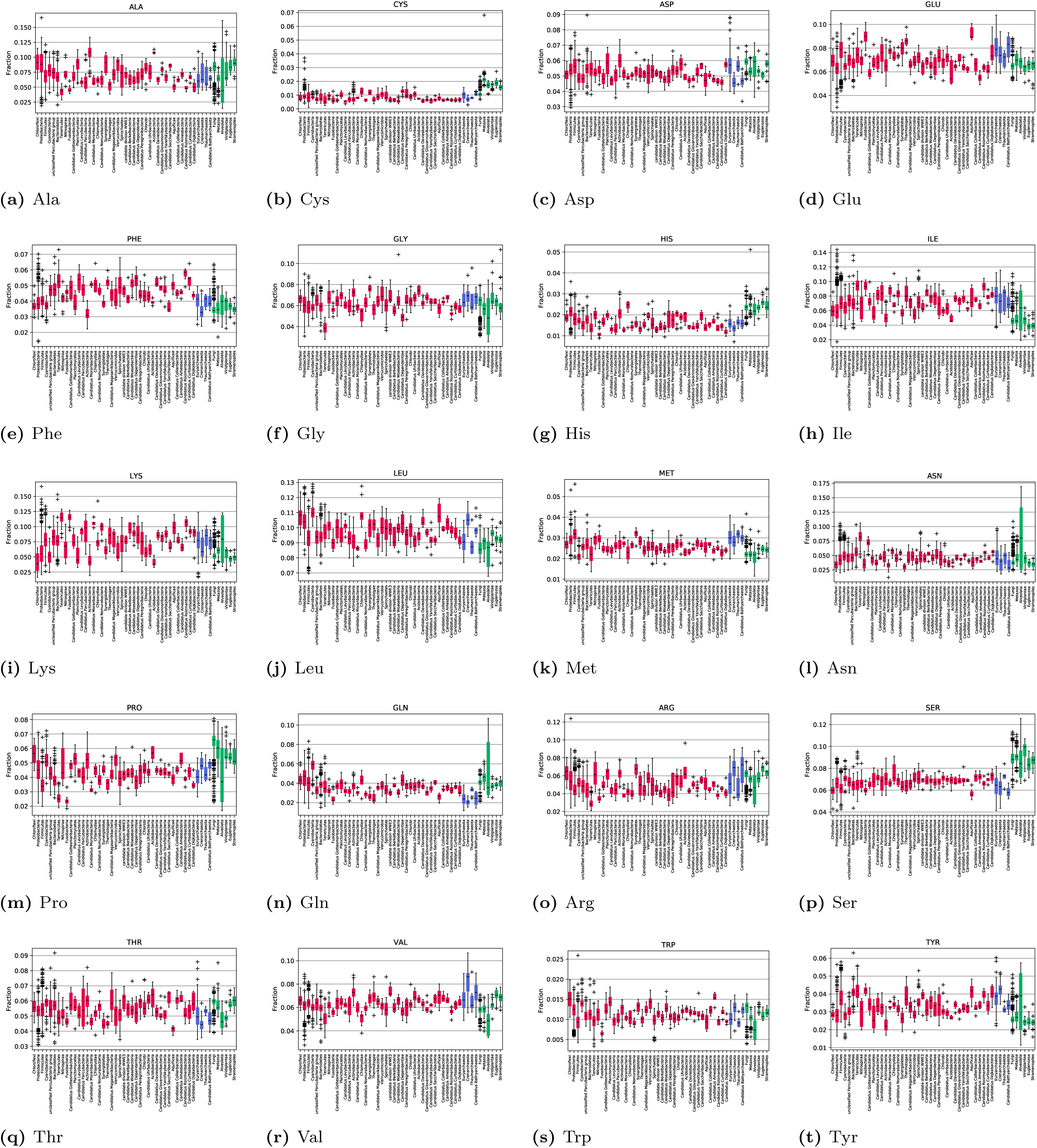
Frequency of amino acids in linker regions grouped by phylum. Bacterial groups are red, eukaryotic green and archaeal blue.

**Figure S5.**
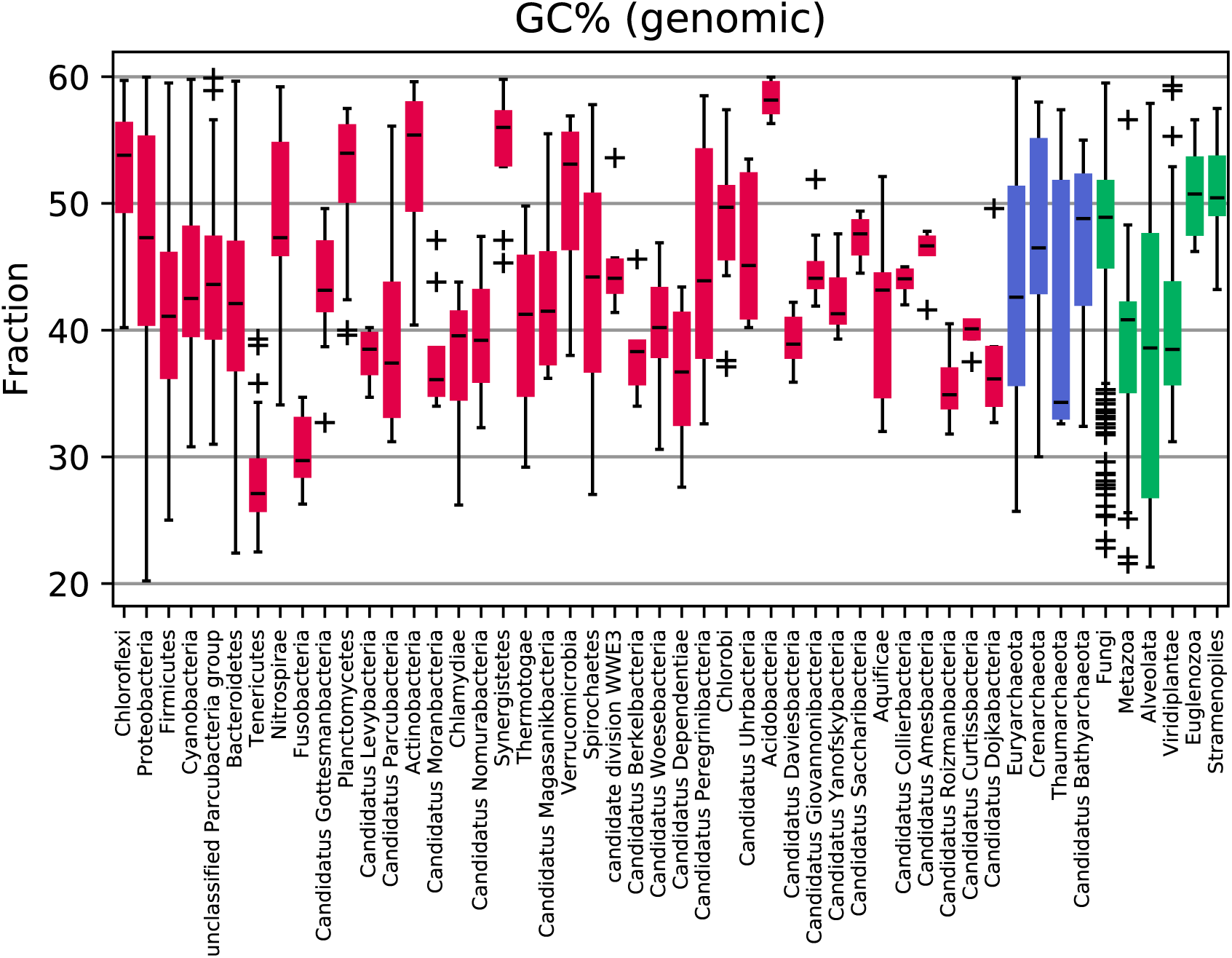
Distribution of GC over different phylogenetic groups (after filtering). Red is bacteria, blue archaea.

**Figure S6.**
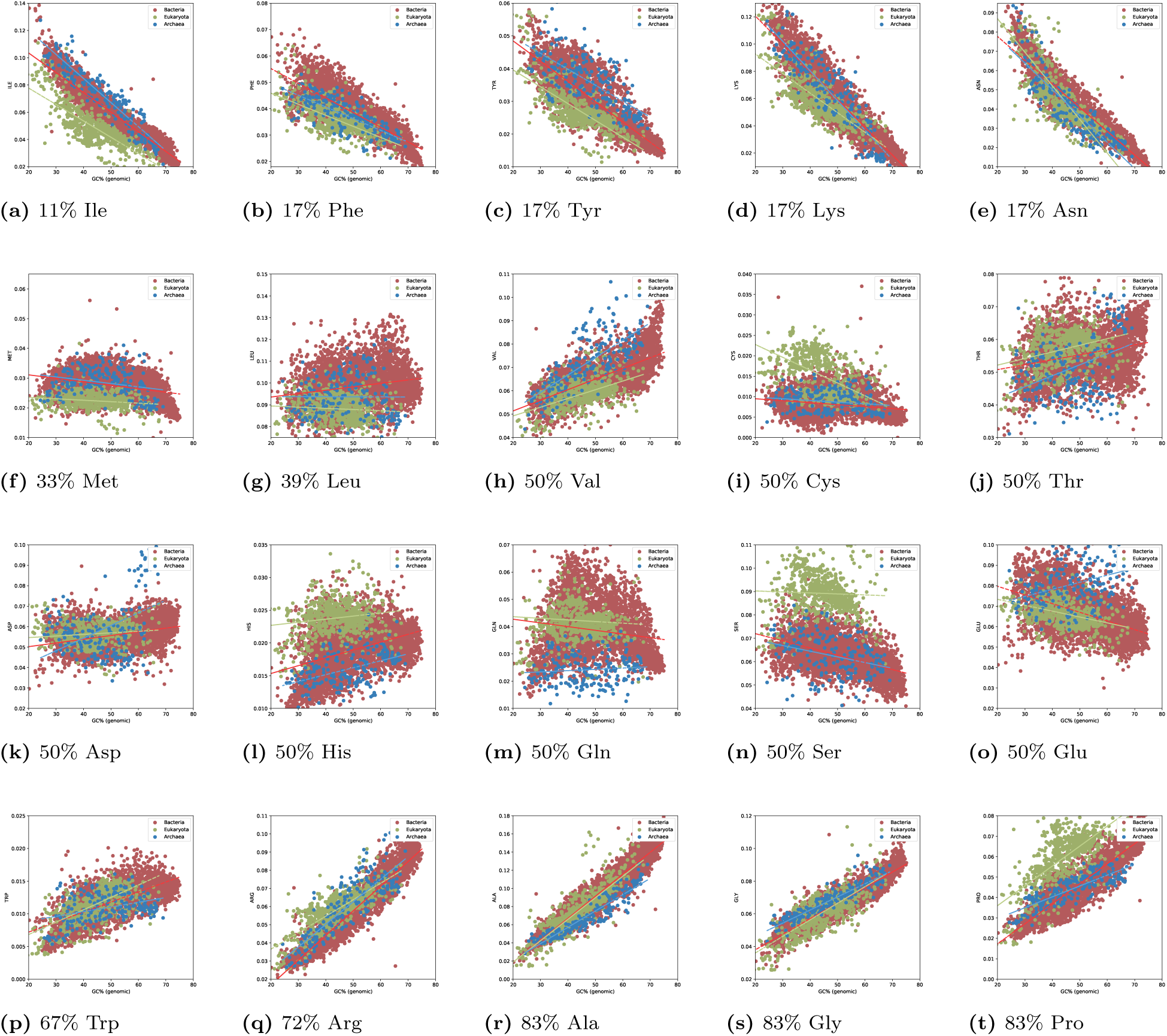
Frequency of all amino acids vs. GC of the linker regions in the genomes. The amino acids are sorted after the GC content of their codons. The number represents the fraction of GC among the codons. The black line represents the expected frequency from codon usage only.

